# Identification, characterization of Apyrase (*APY*) gene family in rice (*Oryza sativa*) and analysis of the expression pattern under various stress conditions

**DOI:** 10.1101/2022.08.12.503798

**Authors:** Aniqua Tasnim Chowdhury, Md. Nazmul Hasan, Fahmid H Bhuiyan, Md. Qamrul Islam, Md. Rakib Wazed Nayon, Md. Mashiur Rahaman, Hammadul Hoque, Nurnabi Azad Jewel, Md Ashrafuzzaman, Shamsul H. Prodhan

**Affiliations:** Department of Genetic Engineering and Biotechnology, School of Life Sciences, Shahjalal University of Science and Technology, Sylhet, Bangladesh; Plant Biotechnology Division, National Institute of Biotechnology, Ganakbari, Ashulia, Savar, Dhaka, Bangladesh; Institute of Epidemiology, Disease Control and Research (IEDCR), Dhaka, Bangladesh

**Keywords:** Rice, Apyrase, Abiotic and biotic stress, Conserved motif, Protein structure, Cis- regulatory elements, Expression pattern.

## Abstract

Apyrase (*APY*) is a nucleoside triphosphate (NTP) diphosphohydrolase (NTPDase) which is a member of the superfamily of guanosine diphosphatase 1 (GDA1) - cluster of differentiation 39 (CD39) nucleoside phosphatase. Under various circumstances like stress, cell growth, the extracellular adenosine triphosphate (eATP) level increases, causing a detrimental influence on cells such as cell growth retardation, ROS production, NO burst, and apoptosis. Apyrase hydrolyses eATP accumulated in the extracellular membrane during stress, wounds, into adenosine diphosphate (ADP) and adenosine monophosphate (AMP) and regulates the stress-responsive pathway in plants. This study was designed for the identification, characterization, and for analysis of *APY* gene expression in *Oryza sativa*. This investigation discovered nine *APY*s in rice, including both endo- and ecto-apyrase. According to duplication event analysis, in the evolution of *OsAPY*s, a significant role is performed by segmental duplication. Their role in stress control, hormonal responsiveness, and the development of cells is supported by the corresponding cis-elements present in their promoter regions. According to expression profiling by RNA-seq data, the genes were expressed in various tissues. Upon exposure to a variety of biotic as well as abiotic stimuli, including anoxia, drought, submergence, alkali, heat, dehydration, salt, and cold, they showed a differential expression pattern. The expression analysis from the RT-qPCR data also showed expression under various abiotic stress conditions, comprising cold, salinity, cadmium, drought, submergence, and especially heat stress. This finding will pave the way for future *in-vivo* analysis, unveil the molecular mechanisms of *APY* genes in stress response, and contribute to the development of stress-tolerant rice varieties.

## Introduction

Apyrase (*APY*), which is a class of nucleoside triphosphate (NTP) diphosphohydrolase (NTPDase) and a member of the superfamily of GDA1-CD39 (guanosine diphosphatase 1- cluster of differentiation 39) nucleoside phosphatase [1], mediates the concentrations of NTP (nucleoside triphosphate) and NDP (nucleoside diphosphate) [2], especially of ATP (adenosine triphosphate) and ADP (adenosine diphosphate) and catalyzes their breakdown and converts them into ADP and AMP (adenosine monophosphate) [3, 4]. ATP, the pervasive intracellular energy resource in every cell, functions in a stress-dependent fashion as it promotes cell growth and development when present in lower concentrations. When the plant cells encounter wound [5], pathogen elicitors ([6, 7], cell growth [8], abiotic stress ([9, 10], touch [9, 11], they discharge ATP into the extracellular matrix (ECM) where it becomes known as eATP (extracellular ATP). Usually, these stresses befall simultaneously and include cross-talk from numerous hormones and signaling pathways [12, 13]. Survival of plants [8], development [4,6,14] gravitropism [15], plant defense strategy [16], cellular apoptosis [17], stress responsiveness [3,5,9,18], are controlled by eATP. Under such circumstances, the eATP level rises, giving rise to many negative impacts on cells like inhibiting the rate of cell growth [19], ROS (reactive oxygen species) production, NO (nitric oxide) burst, as well as triggering apoptosis [20]. At this point, *APY* comes into action by breaking down eATP, which helps plants in cell survival, growth, development, and face off several stresses [21].

The characteristic feature of NTPDase family proteins is to contain five apyrase conserved regions (ACRs) [22], and APYs can be separated as ecto-and endo-APY, as per the subcellular localization [23]. Ecto-APYs are situated on the cell’s surface, and the ones located on cytoplasmic organelles are generally called endo-apyrases. Many apyrase genes include transmembrane domains at their N- and C- terminals [24] and generally feature amino acid glycosylation, which is necessary to ensure proper folding of proteins, targeting at the membrane, cellular distribution, and enzymatic function [1]. However, APYs are not similar to ATPases such as APYs can employ several divalent cofactors like Mn^2+^, Zn^2+^, Ca^2+^, and Mg^2+^, in contrast to ATPases, which employ Mg^2+^ as a cofactor [25]. It also demonstrates insensitivity towards alkaline phosphatase as well as to certain types of ATPase inhibitors [26].

There are several plant species in which *APY*s were discovered, including potato [27, 28], *Arabidopsis* [2], soybean [29], cotton [30], wheat [23]. Especially in *Arabidopsis* and wheat, the gene family has been intensively studied. 7 *APY* members in *Arabidopsis* have been identified and described [2]. AtAPY1 and AtAPY2 are endo-APYs as they are found in the Golgi body. They control various cellular phases [2, 31], and their mutation can dramatically increase extracellular ATP (eATP), which confirms that APYs located intracellularly might also be able to control eATP homeostasis [4, 19]. Others are ecto-APYs, and there have been many pieces of evidence about the involvement of ecto-APYs in the hydrolyzing of eATP [17]. Two other AtAPYs, AtAPY6 and AtAPY7, are also essential to pollen production by controlling the synthesis of polysaccharides [32]. Nine *APY* members were identified in wheat with different expression patterns when subjected to various stresses and even in different tissues [23]. In many species, *APY*s have been found to have involvement in different stress tolerance, such as tolerance towards salinity and drought conditions in *Arabidopsis* after overexpression of *PeAPY1* and *PeAPY2* [33], tolerance to waterlogging in soybean [34]. *PsAPY*, when expressed ectopically, has been reported to exhibit resistance in tobacco to pathogen attack [35]. However, the total signaling pathway working behind the activity of *APY* is still not explained.

Rice is among the world’s most widely grown as well as consumed cereals. The majority of the world relies on rice as a main meal [36]. Agricultural production is at risk due to climate change. Likewise, rice production is being hampered due to climate anomalies, including submergence, less rainfall, cold weather, increasing salinity levels, and many more. Fungal and bacterial blight are two of the most severe and prevalent rice diseases that reduce annual rice production [37]. Promoting stress-resistant rice may increase productivity tremendously as the genome-editing and genetic transfer technologies are improving rapidly; these techniques have created an opportunity to produce stress-resistant crops. With the availability of information on the genomes for several crop species during recent years, a systematic genome-wide study has been readily accessible on stress-associated gene families utilizing bioinformatics techniques. Rice is the first food grain of which the entire genome sequence is available [37]. It provides a chance to locate the genes and systematically categorize the genes and biochemical pathways essential for expanding rice production, conferring tolerance towards several stresses, as well as enhancing the product’s quality.

Different families of genes such as *PYL* [38], *NAC* [39], *GRAS* [40], *MYB* [41], *WRKY* [42], *DREB* [43] were reported in rice as gene families associated with responsiveness towards stress. Many genes have been reported to show specific responses when subjected to different stress conditions. For example, cold, high salinity, and drought are associated with the induction of *OsNAC6/SNAC2* [44]. During dehydration, the HKT-1 protein is required for cell osmoregulation and the maintenance of turgor [45]; the HSP20 family provides tolerance to prolonged heat stress and resistance to different environmental stresses [44]. The CRT/DREBP protein family is mainly regulated by cold stress; on the other hand, DREBP (DRE-binding protein) family induction is caused by drought and high salt stress [46]. Overexpression of different genes responsive to stress in rice could increase the tolerance level when subjected to such stress. Such genes include *OsNHX1* in salt stress [47], *OsDREB* under drought conditions [48], *Sub1A* when subjected to submergence stress [49]. Identifying and characterizing such genes responsive to stresses and applying molecular methods could pave the way toward developing rice varieties tolerant of several stresses.

Since there is ample evidence about the role of *APY*s during the stress response, their identifying and characterizing in plants might help find new development sites for strengthening responsiveness towards several stresses through different gene modification methods. There is an absence of studies about *APY*s in rice. The purpose of this research was to characterize *APY* genes in *O. sativa* and provide insight into their function throughout development and in response to stress by doing a genome-wide identification and analyzing their expression profiles. Rice *APY* family members were identified using bioinformatics techniques throughout this study. We further studied the functional characterization, including phylogenetic relationship analysis with the *APY*s from *Arabidopsis* and wheat. Both RT-qPCR and RNA-seq data were utilized in profiling the pattern of gene expression of the *OsAPY*s and suggested their importance in the process of development as well as response to stress. Conclusively, this result will provide a foothold in further analyzing the roles of *OsAPY* in different stress conditions and in identifying targets to enhance rice’s tolerance level under various stresses.

## Materials and methods

### Identification of *APY* family members in *Oryza sativa*

The TAIR database (https://www.arabidopsis.org/) [50] was employed to get the sequences of the proteins for the seven genes (At3g04080, At5g18280, At1g14240, At1g14230, At1g14250, At2g02970, and At4g19180) of *Arabidopsis thalian*a that belong to the apyrase gene family [2]. On Phytozome v12.1 (https://phytozome.jgi.doe.ov) [51], these sequences were used to run a BLASTp search against the *Oryza sativa* v7 jGI (Rice) dataset. They were also used for the BLASTp search against the NCBI protein database (https://www.ncbi.nlm.nih.gov/protein) and the Rice genome annotation project (http://rice.plantbiology.msu.edu/) [52] separately so that no putative member was missed out. The best hits from these databases were subjected to domain analysis via InterPro (http://www.ebi.ac.uk/interpro/) [53] to ensure the presence of the GDA1-CD39 domain, which is a characteristic feature of APYs. Entries having this domain were chosen for multiple sequence alignment, which was conducted via MEGA X (Molecular Evolutionary Genetics Analysis) software [54]. Their multiple sequence alignment was visualized via GeneDoc version 2.7 [55] to search for the presence of 5 ACRs [22]. Sequences that contained these 5 ACRs were chosen as members of the apyrase family. To confirm that all members of the apyrase family were included, those sequences were submitted as input sequences for a BLAST analysis in Phytozome [51]. The coding sequence (CDS), genomic, and the peptide sequence of the *OsAPY*s were obtained from Phytozome [51].

### Analysis of the physicochemical properties

For assessment of the physicochemical characteristics of the genes, the theoretical isoelectric point (pI), index of instability, grand average of hydropathy (GRAVY), as well as the molecular weight of the proteins were determined. It was done via ProtParam hosted by ExPASy (https://web.expasy.org/protparam/) [56] by uploading the protein sequence. By uploading the protein sequences to WoLF PSORT (https://wolfpsort.hgc.jp/) [57] and CELLO v.2.5 (http://cello.life.nctu.edu.tw/) [58], the subcellular distribution of the proteins was determined.

### Analysis of conserved motifs and gene structure

MEME version 5.3.0 (Multiple Em for Motif Elicitation; http://meme-suite.org) [59] was employed to investigate the conserved motifs on proteins in classic mode, and the motif number was determined to be 10. The minimum width was kept to 6, and the maximum one was 50; each motif’s highest and lowest sites were set to 600 and 2 accordingly. Depending on the appearance frequency of the motifs in MEME [59], they were numbered sequentially and positioned beside their corresponding OsAPYs according to their phylogenetic groups. In order to determine their role, each of the ten detected motifs was evaluated via Pfam (http://pfam.xfam.org/) [60].

*OsAPY* CDS sequences were compared against their respective genomic sequences lacking the UTR (untranslated region) to assess gene structure. Gene structure was determined using the GSDS 2.0 (Gene Structure Display Server) tool (http://gsds.gao-lab.org/) [61], and this web server graphically depicted the gene structure of *OsAPY* genes.

### Analysis of phylogenetic relationship

TAIR [50] along with EnsemblPlants (https://plants.ensembl.org) [62] database was employed to get the peptide sequences of *APY*s from *Arabidopsis thaliana* and *Triticum aestivum*, correspondingly. An unrooted phylogenetic tree was created utilizing MEGA X [54]. It was generated using protein sequences from *Oryza sativa*, *Arabidopsis thaliana*, and *Triticum aestivum*. The Jones-Taylor-Thornton (JTT) substitution model was applied with 1000 bootstrap replications using the maximum likelihood method [63] to keep the rates uniform across sites. It was visualized using the web tool iTOL (https://itol.embl.de/) [64].

### Prediction of gene duplication

PGDD (Plant genome duplication database) (http://chibba.agtec.uga.edu/duplication/) [65] was employed to examine the gene duplication events and to find out the orthologous and paralogous gene pairs. PGDD [65] provided the Ka (synonymous substitution rate) and Ks (non-synonymous substitution rate) values for the duplicated gene pairs. Using the Ks values and a clockwise rate of synonymous substitution (λ) of 1.5 x 10^-8^, the approximate timing of duplication was computed following the formula T = Ks/2λ [66]. A circle plot of the paralogous genes was created using TBTools (https://github.com/CJ-Chen/TBtools/releases) [67].

### Study of the cis-regulatory elements

The sequence located 1000 bp upstream of *OsAPY*s was retrieved from the genomic sequence of rice using Phytozome [51] for investigating the cis-elements and their function. PlantCARE (http://bioinformatics.psb.ugent.be/webtools/plantcare/htmL/) [68] was utilized for predicting the cis-elements. Cis elements were categorized into different groups, and their functions were identified from the literature review. TBTools [67] was utilized to create a visual description of the cis-regulatory elements.

### Analysis of chromosomal distribution

Information from rice genome sequences was used to build a chromosomal organization of *OsAPY* genes based on their location. Phytozome [51] provided details on the number of chromosomes, gene loci, as well as length for the physical map. MapChart (http://www.joinmap.nl) [69] software was used to create a rudimentary physical map of the *OsAPY* gene family, illustrating its location and distribution.

### Specification of *OsAPY* targeting miRNAs

For the identification of miRNAs which target *APY* genes in rice, psRNATarget (http://plantgrn.noble.org/psRNATarget/) [70] was used, and it was tested against every mature rice miRNAs present in the database of miRbase [71]. Using Cytoscape (https://cytoscape.org/) [72], the predicted miRNAs were deployed to build a network. To perform the analysis of pre- computed expression of the detected miRNAs, miRid was employed to search against the rice datasets of the PmiRExAt (http://pmirexat.nabi.res.in/) [73] and a heatmap was generated via GraphPad Prism 9.0.0 (https://www.graphpad.com/company/) [74].

### Identification of SNPs in *APY* genes

The sequences of *APY* genes from 11 rice varieties with different genotypes based on responses towards abiotic stress [38, 75] were compared to the reference Nipponbare sequence using the Rice SNP-Seek database (https://snp-seek.irri.org/index.zul) [76]. SNPs that could be predominantly attributed to variations in the peptide sequence were recognized throughout the genotypes and were termed single amino acid polymorphisms (SAPs).

### Analysis of secondary and tertiary structure of proteins

The secondary structures were generated utilizing the STRIDE program (http://webclu.bio.wzw.tum.de/stride/) [77]. The number and percentage of α helices, extended beta-sheet, turns, coils, and 310-helix were displayed in various colors as part of the structural study. The membrane-spanning motif was investigated by using default parameters of the online tool TMHMM server v.2.0 (http://www.cbs.dtu.dk/services/TMHMM/) [78].

The tertiary structure was generated using the RoseTTAFold method of the Robetta server (https://robetta.bakerlab.org/) [79]. Refinement of the created structures was done via the GalaxyRefine web server (http://galaxy.seoklab.org/refine) [80], and then the energy minimization was done using Swiss-PdbViewer v4.1 (http://www.expasy.org/spdbv/) [81]. PROCHECK (https://servicesn.mbi.ucla.edu/PROCHECK/) [82] and ERRAT (https://servicesn.mbi.ucla.edu/ERRAT/) [83] servers were further used for the validation of the structure. For the assessment of Z-scores and energy plots, ProSA-web (https://prosa.services.came.sbg.ac.at/prosa.php) [84] was utilized. Discovery Studio software was finally used to visualize the tertiary protein structures [85].

### Analysis of protein-ligand docking

Molecular docking is an *in-silico* strategy for assessing the binding affinity of a ligand to a receptor molecule. Since ATP is a vital ligand of APY proteins, it was chosen as the ligand for the study. HDOCK server (http://hdock.phys.hust.edu.cn/) [86] was used as the docking tool to conduct the docking analysis between the OsAPY proteins and ATP. It was done using the default parameters. The 3D structure of ATP was downloaded from the PubChem database (PubChem CID: 5957). After docking, the top 10 solution complex generated via HDOCK [86] were downloaded, and their structures were further analyzed and validated using PDBsum (http://www.biochem.ucl.ac.uk/bsm/pdbsum) [87], PROCHECK [82] and ERRAT [83] servers. The best structure was selected, and it was aligned and visualized. Then finally, the protein and ligand residue interaction was studied using Discovery Studio [85].

### Study of expression profiling of *OsAPY*s in rice using RNA-seq data

Rice Genome Annotation Project [52] was utilized to collect the data for the expression of *APY* genes under several tissues. Accordingly, the RNA-seq expression values were listed. GraphPad Prism 9.0.0 [74] software was used to create a heat map from the data. GENEVESTIGATOR (https://genevestigator.com/gv/) [88] provided the pattern of *OsAPY* expression in various stresses, and the log2 values of relative expression were used to generate the heat map via GraphPad Prism 9.0.0 [74].

### Plant germination, treatment, and total RNA isolation

The healthy, high-quality and mature seeds of BRRI dhan28 were obtained from the Bangladesh Rice Research Institute (BRRI) and used in this experiment. They were washed thoroughly and then were allowed to germinate on a petri dish containing water-soaked tissue paper. After 3-4 days, the germinated seedlings were transferred to the hydroponic culture system. A controlled condition was maintained in the culture room, including a temperature of 25±2 °C with a photoperiod consisting of 16 hours of light followed by 8 hours of darkness and 1500-2000 lux light intensity. After 20 days, the seedlings were subjected to different stresses for 18-20 hours which included cadmium stress (100mM CdCl_2_ dissolved in distilled water), cold (4 °C), salinity (100mM NaCl dissolved in distilled water), submergence, drought (3 mg/L polyethylene glycol dissolved in distilled water), heat stress (42 °C). As control, seedlings that had been untreated were utilized. After the treatment, fresh young leaves were collected and washed several times with 70% alcohol and distilled water in order to extract the RNA. Using Invitrogen^TM^ TriZOL^TM^ reagent (Thermo Fisher Scientific Corporation, USA), total RNA was isolated from leaves. The extracted RNA was subsequently processed with DNase I of Invitrogen^TM^ DNA-free^TM^ DNA Removal Kit (Thermo Fisher Scientific Corporation, USA) to remove the genomic DNA contamination. Finally, the GoScript^TM^ Reverse Transcription System (Promega Corporation, USA) was utilized to synthesize the complementary DNA (cDNA). These procedures were carried out per the protocol of the manufacturers.

### Analysis of gene expression under abiotic stress conditions using RT-qPCR data

Primer3 v.0.4.0 (https://bioinfo.ut.ee/primer3-0.4.0/) [89] was used to generate primers for the RT-qPCR, keeping the product length between 208 and 220 bp. The real-time PCR was done using GoTaq® qPCR Master Mix (2X) (Promega Corporation, USA) on CFX96^TM^ Real-Time PCR Detection System (BioRad). For the reference gene, eukaryotic elongation factor 1 alpha (*eEF-1α*) was chosen [90]. Each set of gene-specific primers took up 1 μL of the 15 μL reaction mixture, which also comprised 7.5 μL of GoTaq® qPCR Master Mix (2X), 2 μL of cDNA (10 times diluted), and 3.5 μL of nuclease-free water. In order to carry out each of the reactions, the following conditions were applied: at 95 °C initial denaturation for 10 min preceding 40 cycles of denaturation at 95 °C for 15s, annealing for 30s, and extension at 72° C for 40s. The annealing temperature was 55.4 °C for *OsAPY2* and *OsAPY4*; 56.4 °C for *OsAPY1* and *OsAPY5*; 57.4 °C for *OsAPY3*, *OsAPY6*, and *OsAPY8*; 58.4 °C for *OsAPY7* and *OsAPY9*; 55.7 °C for *eEF-1α*. Three separate experiments were carried out for each sample and the melting curve analysis was done after the PCR amplification. The delta-delta Ct value approach [91] was utilized in calculating the ratio of relative expression. Technical replication was used to find the mean value of expression at different treatments. MS Office 365 and GraphPad Prism

9.0.0 [74] were used to analyze the data. A one-way ANOVA, followed by a Bonferroni post hoc test, was used to assess the significant differences (P ≤ 0.05). To represent the significant differences, different means were labeled with different number of stars (*).

## Results

### Identification of *APY* family members in rice and study of their physicochemical properties

Rice was found to contain nine members of the *APY* gene family. To BLAST against the rice genome, the *Arabidopsis APY*s were utilized as reference sequences. Upon performing BLASTp searches against the *Oryza sativa*, a total of 19 hits were found containing the GDA1- CD39 domain. All the domain annotations were confirmed by sequence analysis in InterPro [53].

These 19 transcripts belonged to a total of 12 genes. The primary transcripts annotated by Phytozome [51] were selected as the representative transcripts of the genes with alternative splice forms. 9 of the 12 genes were found to possess all five Apyrase Conserved Regions (ACRs) (S1 Fig); hence they were designated Apyrase genes. For the naming of the genes (*OsAPY1*-*OsAPY9*), the prefix ’Os’ for *Oryza sativa* was used, followed by ’APY’ for Apyrase, and then the sequential number that corresponded to their chromosome number and location was used. Table 1 lists the physicochemical properties of all representatives of the *OsAPY* Family.

**Table 1.** Physicochemical properties of the members of the apyrase gene family in rice (*O. sativa*).

| Gene Name | Locus Name | Length |  |  | MW (Da) | pI | Instability Index | GRAVY Value | Localization |
| --- | --- | --- | --- | --- | --- | --- | --- | --- | --- |
|  |  | Genomic (bp) | CDS (bp) | Protein (aa) |  |  |  |  |  |
| *OsAPY1* | LOC_Os03g21120 | 5548 | 1470 | 489 | 52799.01 | 5.72 | 39.76<br>(Stable) | -0.09 | Cp<sup>ab</sup> |
| *OsAPY2* | LOC_Os03g26080 | 3847 | 1527 | 508 | 55346.06 | 9.1 | 53.33<br>(Unstable) | -0.204 | PM<sup>ab</sup> |
| *OsAPY3* | LOC_Os07g48430 | 4260 | 1404 | 467 | 50437.74 | 8.97 | 36.76<br>(Stable) | -0.136 | Mt<sup>a</sup>, Cp<sup>b</sup> |
| *OsAPY4* | LOC_Os08g33850 | 4217 | 1629 | 542 | 59596.12 | 9.34 | 48.76<br>(Unstable) | -0.149 | PM<sup>a</sup>, ER<sup>b</sup> |
| *OsAPY5* | LOC_Os10g21000 | 4624 | 2109 | 702 | 77242.54 | 9.18 | 43.81<br>(Unstable) | -0.171 | PM<sup>ab</sup> |
| *OsAPY6* | LOC_Os11g03270 | 6059 | 1647 | 548 | 59542.4 | 7.96 | 36.68<br>(Stable) | -0.077 | Cp<sup>ab</sup> |
| *OsAPY7* | LOC_Os11g03290 | 3902 | 1374 | 457 | 49622.22 | 5.44 | 31.59<br>(Stable) | -0.179 | Cp<sup>ab</sup> |
| *OsAPY8* | LOC_Os11g25260 | 7876 | 1428 | 475 | 50321.86 | 5.83 | 37.31<br>(Stable) | -0.12 | Cp<sup>ab</sup> |
| *OsAPY9* | LOC_Os12g02980 | 3138 | 1356 | 451 | 48900.8 | 8.01 | 30.98<br>(Stable) | -0.155 | Cp<sup>a</sup>, ER<sup>b</sup> |

CDS- Coding sequence, MW- Molecular weight, pI- Isoelectric point, GRAVY- Grand average of hydropathy, Cp- Chloroplast. PM- Plasma membrane, Mt- Mitochondria, ER- Endoplasmic reticulum.
<sup>a</sup>Subcellular localization according to CELLO, <sup>b</sup>subcellular localization according to WoLF PSORT.

The theoretical pI of proteins belonging to the apyrase gene family ranged between 5.44 and 9.34, with an average of 7.73. *OsAPY*4 had the highest and *OsAPY*7 had the lowest pI value. The proteins had molecular weights ranging from 48900.8 Da to 77242.54 Da belonging to *OsAPY*9 and *OsAPY*5, respectively. The average molecular weight of the proteins was 55978.8 Da. The GRAVY value for all the proteins was negative, and as per the data, all protein molecules are hydrophilic [92]. Six genes had an instability index lower than 40, meaning they are stable. It revealed that 67% of the genes are stable, and 33% of them are unstable.

For the prediction of subcellular location, two different tools were used, and per the subcellular localization, both ecto- and endo-APYs are present in this gene family [23]. Their results varied in OsAPY3, OsAPY4, and OsAPY9. According to CELLO [58], they are located on mitochondria, plasma membrane, and chloroplast. On the other hand, according to WoLF PSORT [57], their subcellular localization is on chloroplast for OsAPY3 and endoplasmic reticulum for OsAPY4 and OsAPY9. The predicted subcellular localization of the proteins according to two different tools is given in Table 1.

### Analysis of conserved motifs and gene structure

In total, ten conserved motifs were discovered, which were numbered 1-10 (Fig 1b). 4 motifs, including motif 1, 5, 8, and 10, were conserved and existed in all the OsAPYs. Members of each phylogenetic group had similar motif distribution, which affirms the classification of groups. All the members of group I in the phylogenetic tree (Fig 1a) had the same motif distribution, containing all the ten motifs. On the other hand, the members of group II in the phylogenetic tree, OsAPY2, and OsAPY4, contained almost similar numbers of motifs except motif 2, which was present in OsAPY4 only. Though only OsAPY5 belonged to group III, its motif distribution was the same as OsAPY4 despite being in different phylogenetic groups. Functions of these motifs were also analyzed, but there was no information on the function of motif 9 and 10. Their analysis revealed that the remaining eight motifs belong to the GDA1/CD39 (nucleoside phosphatase) family (S1 Table).

**Fig 1.**
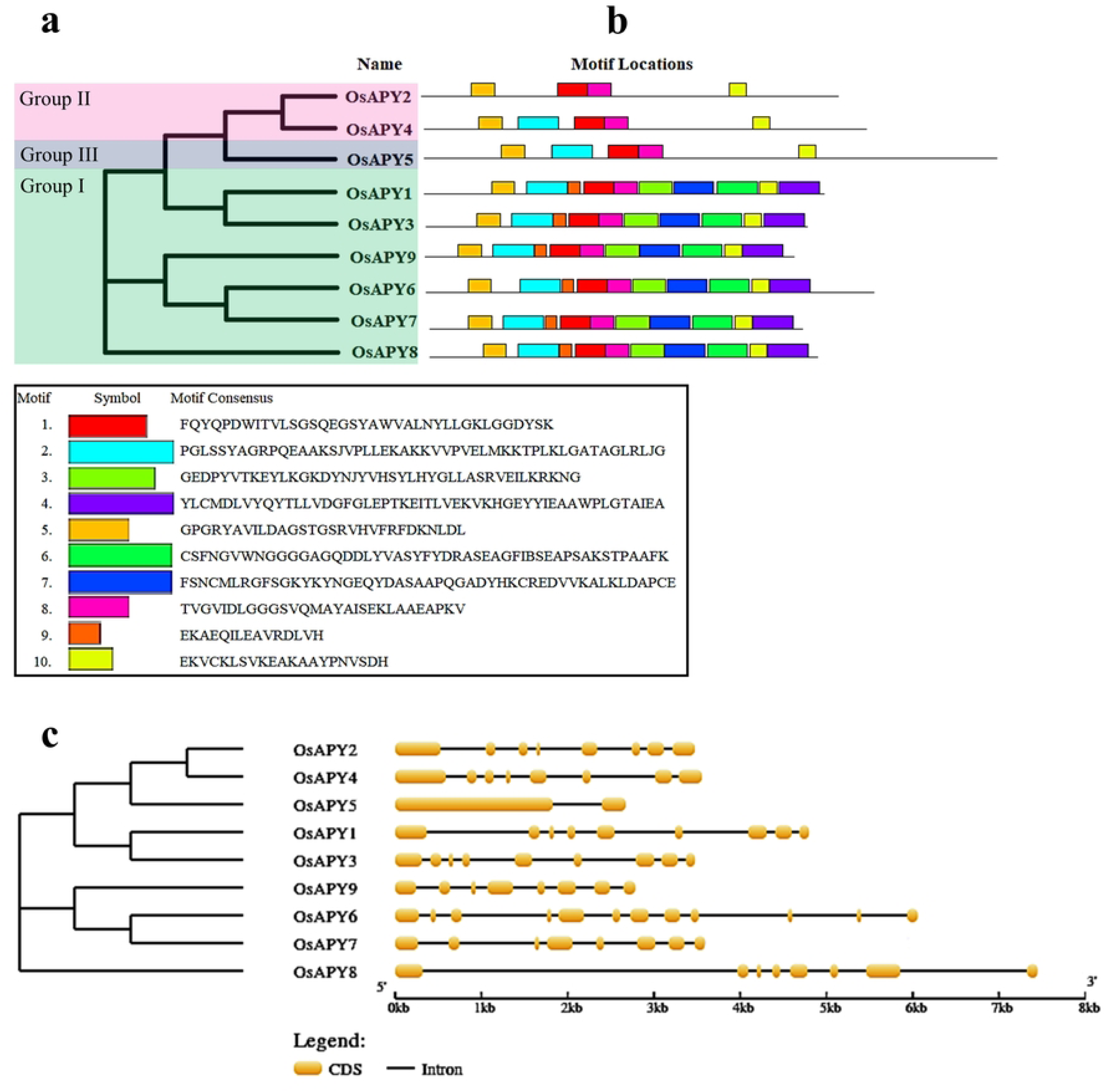
Schematic depiction of phylogenetic relationship, gene structure, and conserved motifs of *APY* genes. a- Phylogenetic relationship among *OsAPY*s generated via maximum likelihood approach. The green, pink and blue boxes represent groups I, II, and III, respectively. b- OsAPY conserved motif distribution. Ten varying-colored boxes represent the motifs. The legend below shows the corresponding motif’s protein sequence. c- Gene structure of *APY*s. Yellow color boxes specify exons, and introns are depicted by lines. The exon-intron lengths can be calculated following the scale shown below.

The exon and intron arrangement of *OsAPY*s was studied in order to better comprehend their structure. The total number of coding sequences (CDS) or exons and introns of the *OsAPY* genes were 2-12 and 1-11, respectively (Fig 1c), and there was no intronless gene. *OsAPY5* contained the lowest number of exons and introns, which are two exons and one intron, and on the other hand, *OsAPY6* contained the highest number of exons and introns, which is 12 exons and 11 introns.

### Analysis of phylogenetic relationship in *OsAPY* family

The maximum likelihood method was utilized to create an unrooted phylogenetic tree to study the evolutionary relationship between the *APY* genes. It was done using the peptide sequences of seven apyrase homologs from *Arabidopsis thaliana* (*AtAPY*) [2], nine apyrase homologs from *Triticum aestivum* (*TaAPY*) [23] and the nine identified apyrase homologs from *Oryza sativa* (*OsAPY*). It was demonstrated by this phylogenetic tree (Fig 2) that the nine *OsAPY* genes could be split into three separate groups referred to as I, II, and III. There were already three distinct groups for the *AtAPY* and *TaAPY* genes [23]. No *OsAPY* gene was classified into the new group. Among the 9 *OsAPY* genes, *OsAPY1, OsAPY3, OsAPY6*-*OsAPY9* belonged to group I. *TaAPY1*-*TaAPY3*.*4* and *AtAPY1*, *AtAPY2* also were in group I. *OsAPY2*, and *OsAPY4* fell in group II. Group II also contained *TaAPY5*, *TaAPY6*, *AtAPY3*-*AtAPY6*. Group III consisted of *OsAPY5*, *AtAPY7*, and *TaAPY7*.

**Fig 2.**
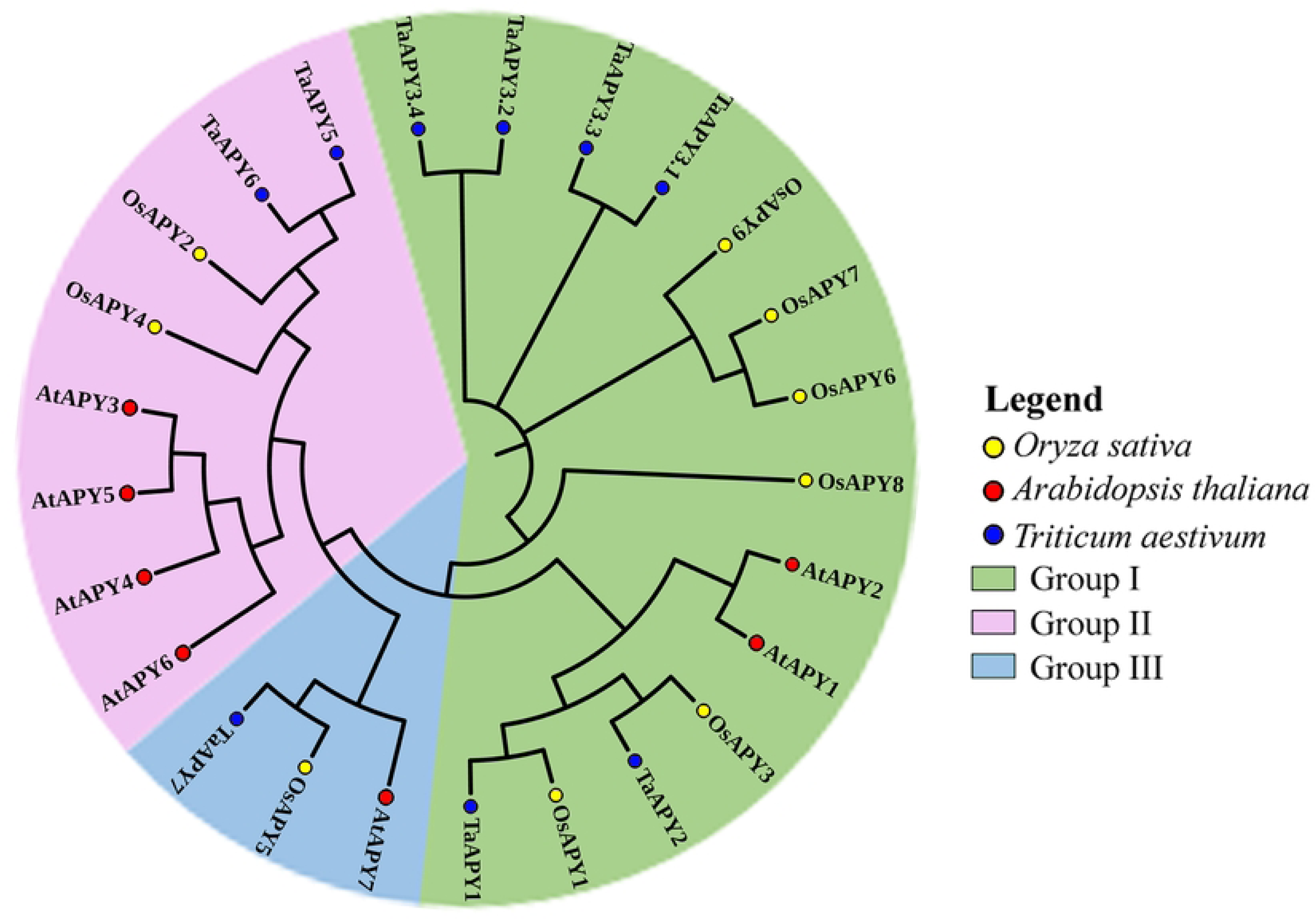
Phylogenetic analysis of the *APY*s in rice, *Arabidopsis*, and wheat. The MEGA X software was employed to generate a phylogenetic tree by utilizing the maximum likelihood approach along with 1000 bootstrap replications. Genes were divided into three different groups, each marked with a different color.

### Analysis of gene duplication

A crucial role is played by duplication events for expanding the gene family and in the emergence of new gene functions. There were six duplications in total in the rice genome, with three occurring within the *APY* family (Fig 3 and S2 Table). Because no genes on the same chromosome had been duplicated, there was no tandem duplication at all. In this study, the duplication type was segmental. The three groups of segmentally duplicated genes within the rice apyrase gene family are *OsAPY1/9*, located on chromosomes 3 and 12, respectively; *OsAPY1/3*, found on chromosomes 3 and 7, respectively; *OsAPY3/9*, situated on chromosomes 7 and 12 respectively.

**Fig 3.**
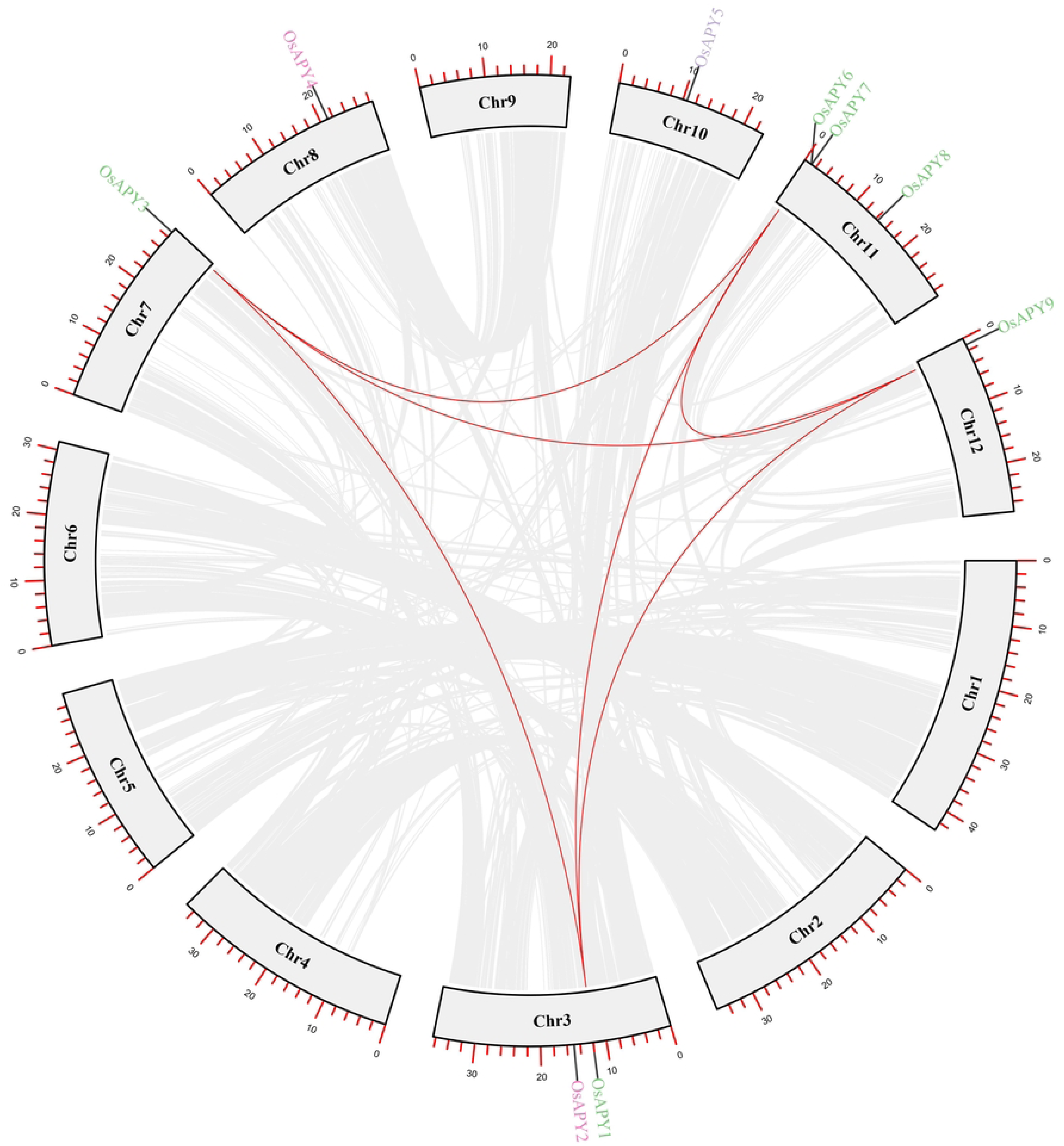
Duplication events of *OsAPY*s within rice genome. Duplication of the *OsAPY*s was specified by the red lines, and the one among the whole rice genome was denoted by the gray lines. TBTools was used to create the circle plot.

The non-synonymous (Ka) and synonymous (Ks) substitution rate of genes was used to measure the selection pressure of duplication occurrences. Ka/Ks = 1 implies neutral selection, whereas Ka/Ks<1 represents purifying selection, and Ka/Ks>1 denotes positive selection [93]. Between 0.1476 to 0.5560, Ka/Ks of segmental duplication had a mean of 0.28021. As demonstrated by Ka/Ks ratios lower than 1, all the genes originated under the influence of purifying selection (S2 Table). The duplication time of each event was counted in million years ago (MYA) unit (S2 Table), and it occurred between 66.6 MYA and 1.54 MYA.

Eight orthologous pairs were found between rice *OsAPY*s and other species (Table 2). It- is duplicated with sorghum, western poplar, cotton, grapevine, maize, and soybean. The range of Ka/Ks was between 0.143 and 0.357, and the mean was 0.2471. These data suggest that duplication events had a significant impact on the expansion and functional diversity, and an important role was served by segmental duplication in the evolution of the *APY* family of genes.

**Table 2.** Duplication event between rice *APY* genes and other species.

| Duplicated Gene 1 | Duplicated Gene 2 | Ka | Ks | Ka/Ks | Duplication time (MYA) | Purifying Selection | Duplication type |
| --- | --- | --- | --- | --- | --- | --- | --- |
| *OsAPY2* | Sobic.001G356200<br>(Sorghum) | 0.1427 | 0.4936 | 0.2891 | 16.45 | Yes | Segmental |
| *OsAPY1* | Potri.019G031000<br>(Western Poplar) | 0.2653 | 1.6902 | 0.157 | 56.34 | Yes | Segmental |
| *OsAPY1* | Gorai.007G268200<br>(Cotton) | 0.3114 | 1.6822 | 0.1851 | 56.07 | Yes | Segmental |
| *OsAPY2* | GRMZM2G056944<br>(Maize) | 0.16 | 0.4481 | 0.3571 | 14.93 | Yes | Segmental |
| *OsAPY2* | GRMZM2G097987<br>(Maize) | 0.158 | 0.4801 | 0.3291 | 16 | Yes | Segmental |
| *OsAPY4* | Glyma.02G107700<br>(Soybean) | 0.4302 | 1.6258 | 0.2646 | 54.19 | Yes | Segmental |
| *OsAPY4* | Glyma.01G047100<br>(Soybean) | 0.4293 | 1.7026 | 0.2521 | 56.75 | Yes | Segmental |
| *OsAPY4* | GSVIVT01019977001<br>(Grape vine) | 0.3133 | 2.1857 | 0.1433 | 72.85 | Yes | Segmental |

### Cis-acting regulatory elements (CAREs) analysis

Rice *APY* genes might be regulated by promoter sequences, which are known to be associated with the regulation of transcription of the genes in plants. Analysis of 1 kbp upstream promoter sequences in *O. sativa* indicated the presence of 69 CAREs. These cis-elements were grouped into eight different functional categories (Fig 4): i) Light responsive cis-elements, ii) Abiotic challenge responsive elements, iii) Hormonal regulation responsive elements, iv) Elements responsible for cellular development, v) Promoter associated elements, vi) Biotic challenge responsive elements, vii) Elements with miscellaneous functions and viii) Elements with unknown functions. Hormonal regulation responsive elements were further divided into a) MeJA responsive, b) Salicylic acid-responsive, c) Auxin responsive, d) Gibberellin responsive, e) Ethylene responsive, and f) Abscisic acid-responsive elements (Fig 4).

**Fig 4.**
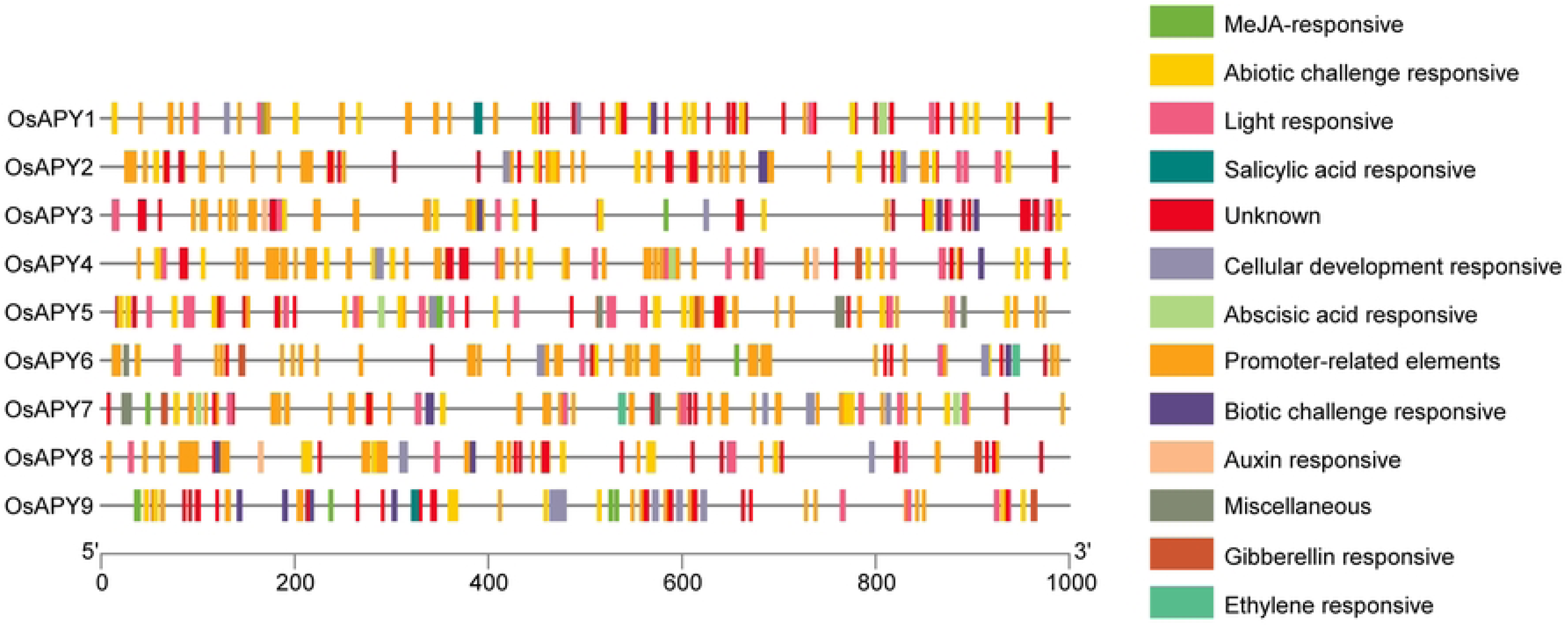
Schematic diagram of predicted cis-elements in *OsAPY* promoter region. Upstream nucleotides of the translation initiation point are shown by the scale at the end. Different colored boxes depict different cis-regulatory elements.

Different abiotic stress-associated elements (anaerobic, anoxic, low temperature, drought, salt, and cold) were observed in the *OsAPY* promoter region (Table 3). STRE, TC-rich repeats, MBS, ARE, GC-motif, LTR, CCAAT-box, AP-1, DRE core, MYB, MYÇ were the abiotic challenge responsive cis-elements. Among all these elements, STRE was present in the highest frequency, and DRE1 was present in the lowest frequency. W box, WRE3, and WUN-motif (Table 3), the biotic challenge responsive elements, mainly act as fungal elicitors and wound-responsive elements. WRE3 occurred in the highest frequency, and WUN-motif occurred in the lowest frequency.

**Table 3.**
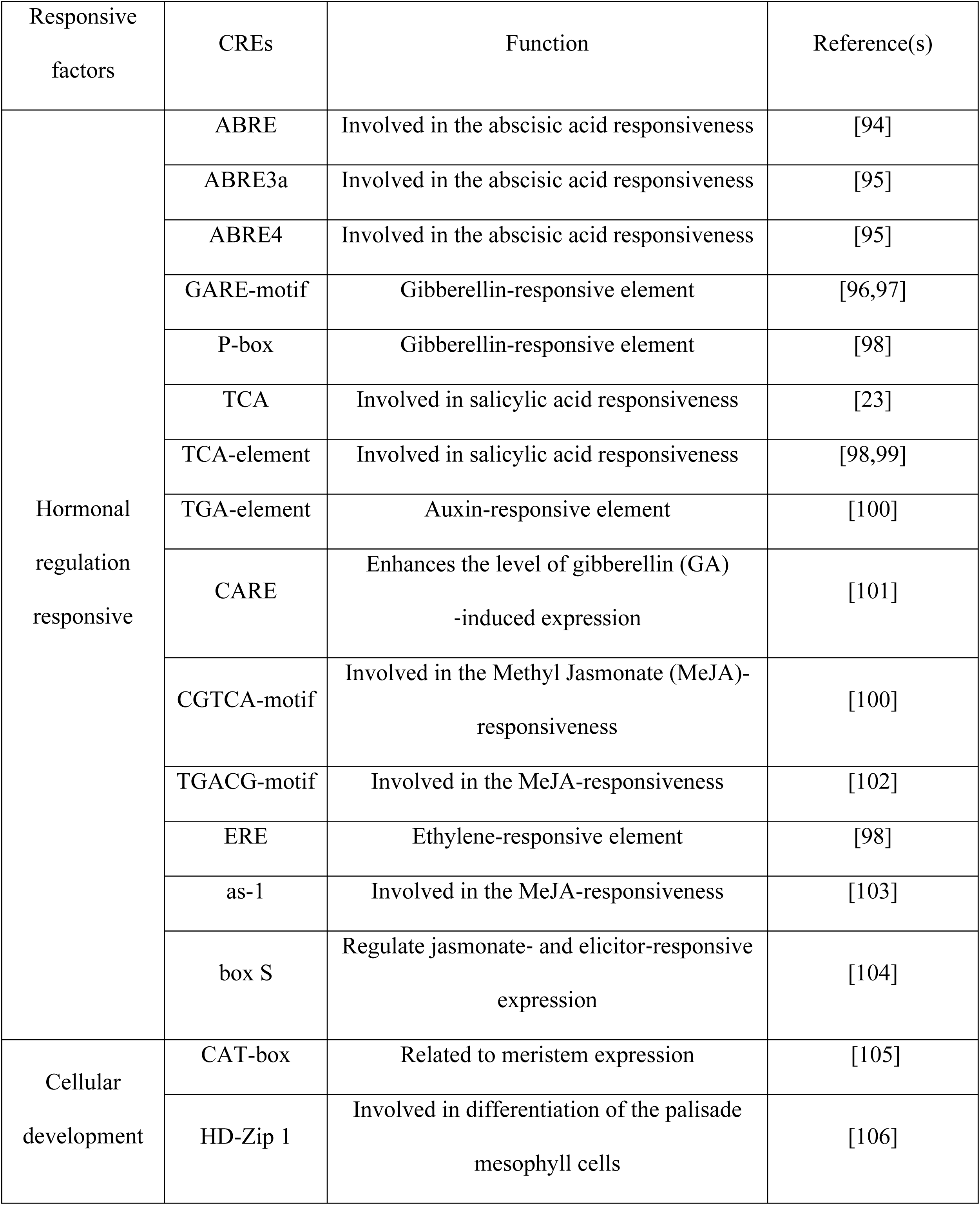
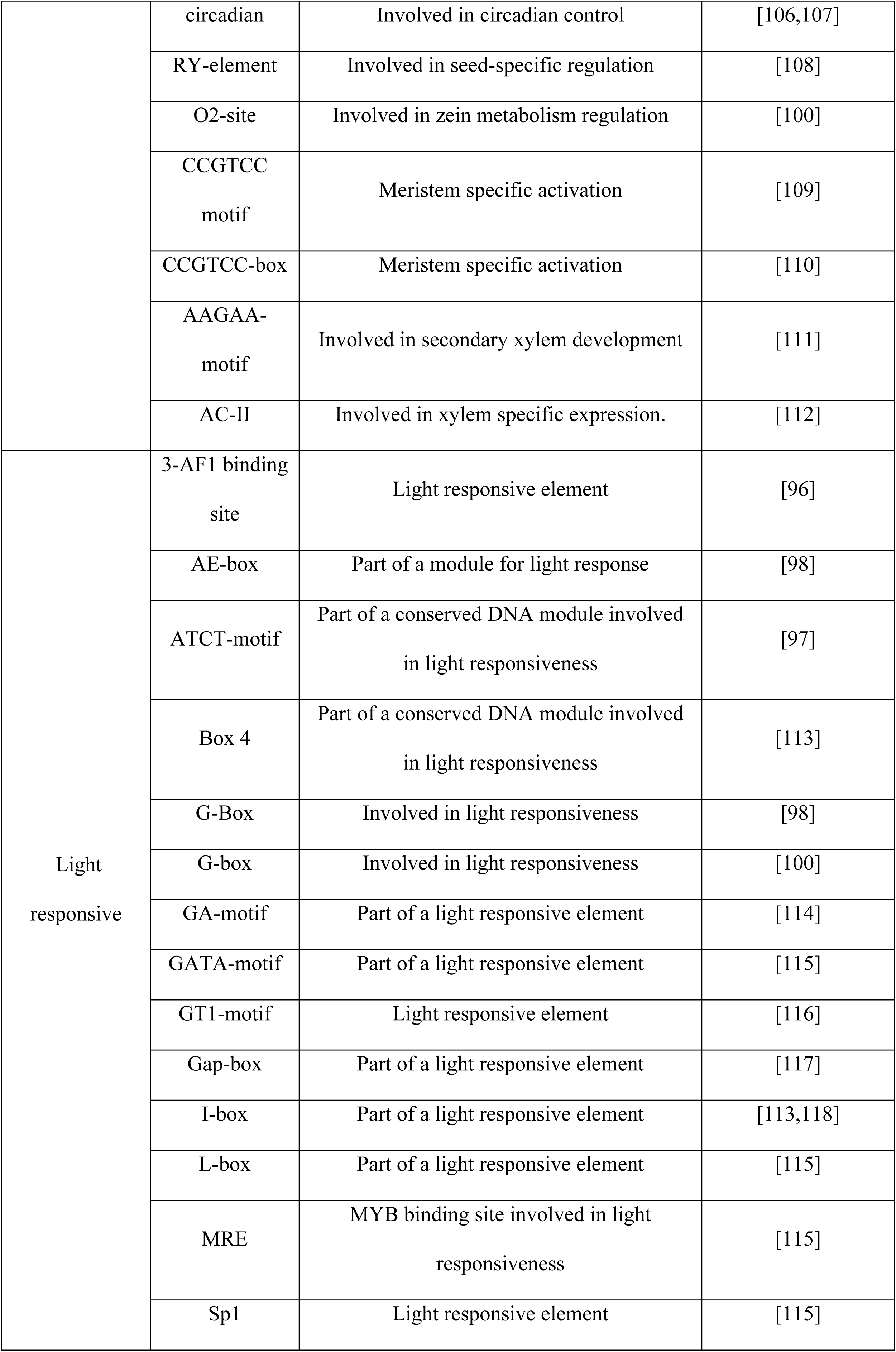
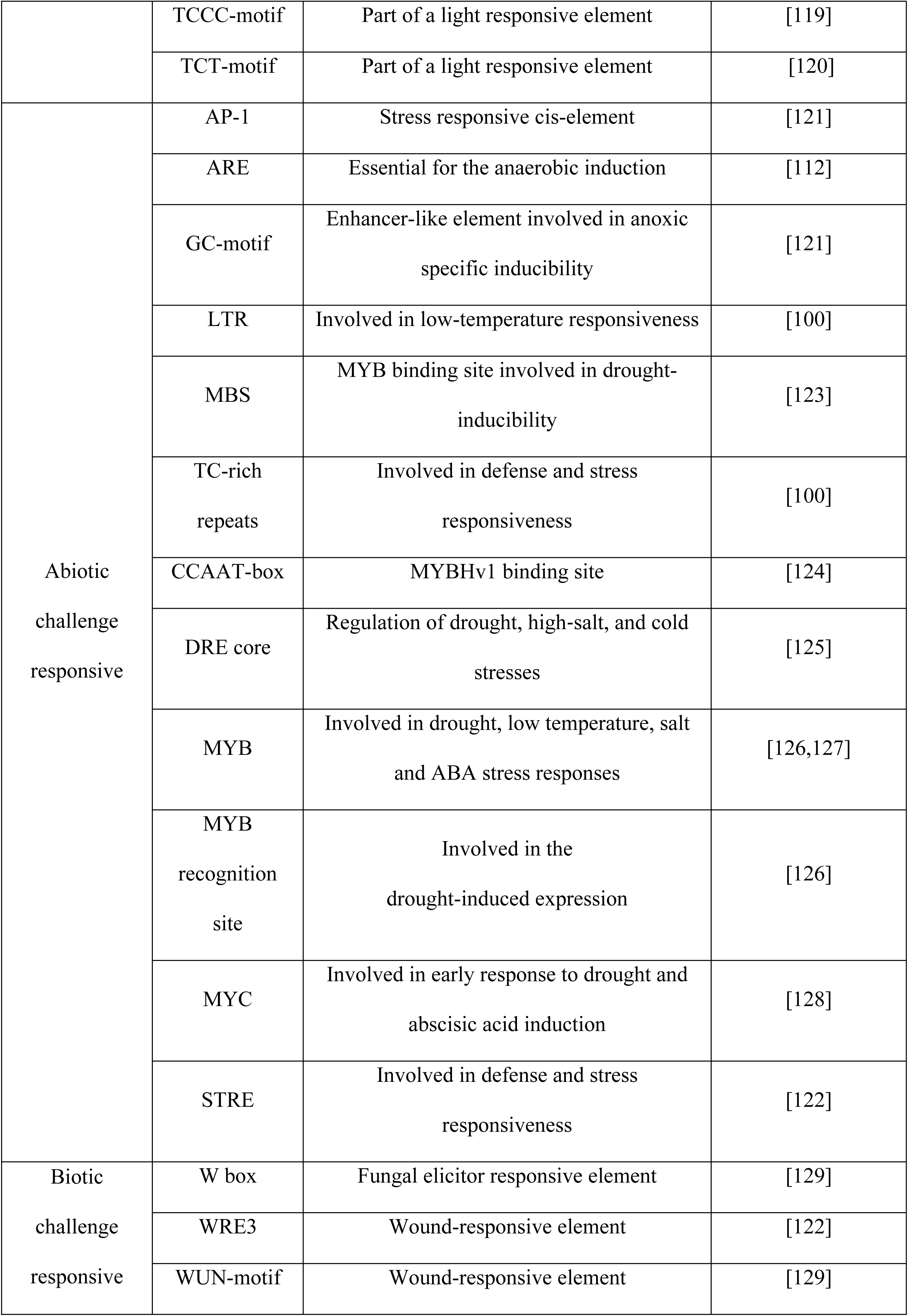
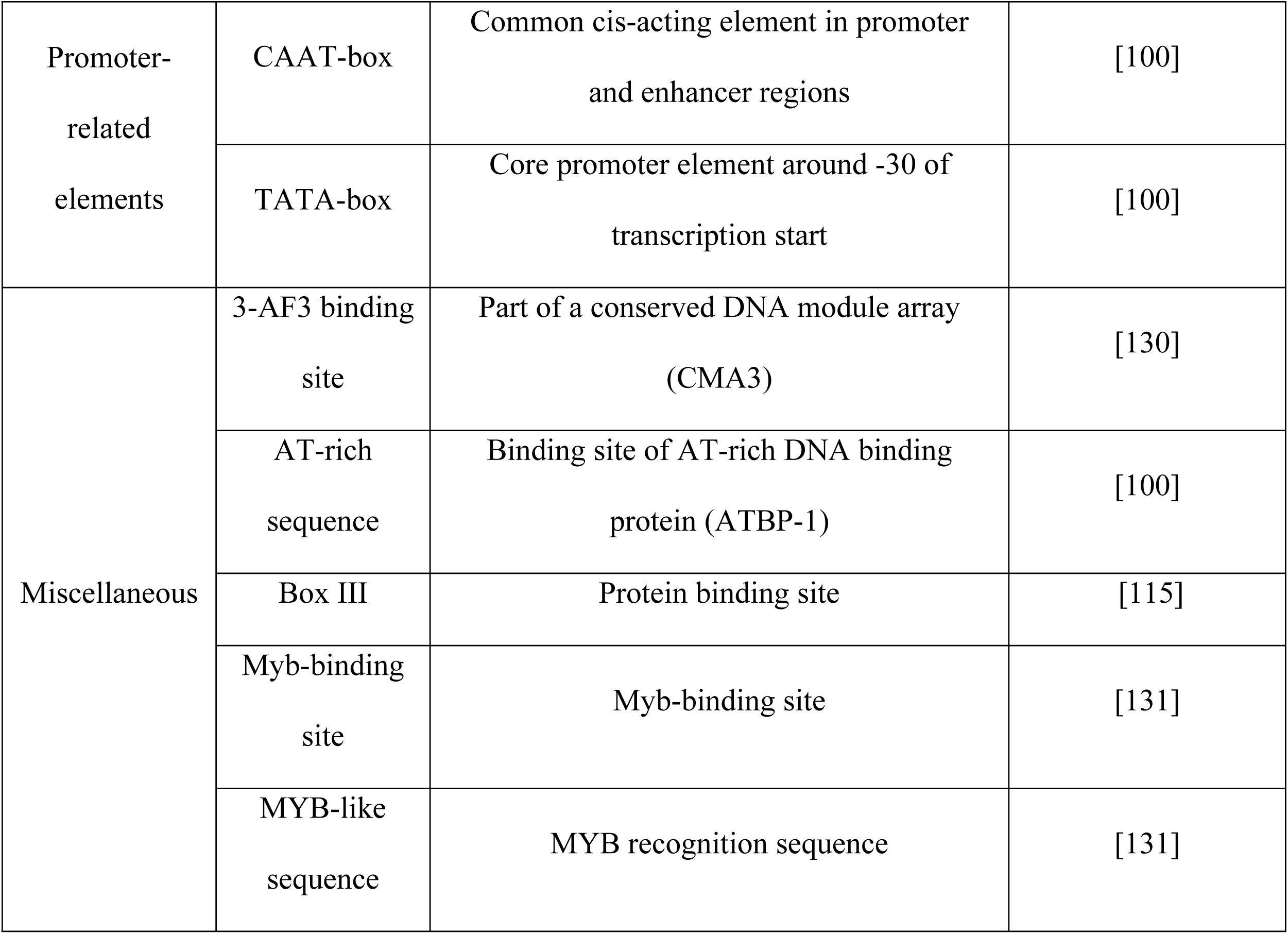
List of cis-regulatory elements (CREs) found in the 5′ UTR region of the *OsAPY*s.

TGACG motif, CGTCA motif, TGA element, TCA element, GARE-motif, ABRE, P-box, ERE, as-1, and box S constituted the hormonal regulation responsive cis-elements (Table 3). Cellular development responsive cis-elements play a major role in the meristem expression, palisade mesophyll tissue differentiation, circadian control, seed-specific regulation, zein metabolism regulation, and xylem-specific expression (Table 3). These cis-elements included CAT-box, HD-Zip 1, circadian, RY-element, O2-site, CCGTCC motif, CCGTCC box, AAGAA-motif, and AC-II.

Light responsive cis-elements included 3-AF1 binding site, TCT-motif, TCCC-motif, MRE, Gap-box, I-box, GT1-motif, ATCT-motif, Box 4, GATA-motif, GA-motif, G-box, G-Box, AE- box, L-box, Sp1, (Table 3). They mainly function as a light responsive element or a module or a component of such an element. TATA-box and CAAT-box were the two elements, related to promoter, found in *OsAPY* genes, and they are core promoter elements that mainly function in promoter and enhancer regions. Cis-elements with miscellaneous functions included 3-AF3 binding site, MYB-like sequence Myb-binding site, Box III, and AT-rich sequence (Table 3). *OsAPY5* contains a 3-AF3 binding site that is a DNA module array component. The AT-rich DNA binding protein (ATBP-1) utilizes a binding site situated on *OsAPY7*, which is located on the AT-rich sequence. On the other hand, Box III is located on *OsAPY7* as a binding site for proteins.

### Analysis of chromosomal distribution

All the 12 chromosomes of rice were examined for *APY* gene distribution, and the results indicated an unequal distribution of *APY* genes (Fig 5). No *OsAPY* member was mapped onto chromosome 1, 2, 4, 5, 6, and 9. It was observed that both *OsAPY1* and *OsAPY2* were on chromosome 3. The chromosomal positions of *OsAPY3*, *4*, *5*, and *9* were 7, 8, 10, and 12, respectively. *OsAPY6-OsAPY8* was located on chromosome 11. *OsAPY1*, *2*, *5*, and *8* were concentrated near the centromere. *OsAPY6*, *7*, and *9* were situated on the p arm of the chromosome, and on the other hand, *OsAPY3* and *4* were placed on the q arm of the chromosome. The location of all the *OsAPY*s in different chromosomes, their position, and orientation are mentioned in the S3 Table.

**Fig 5.**
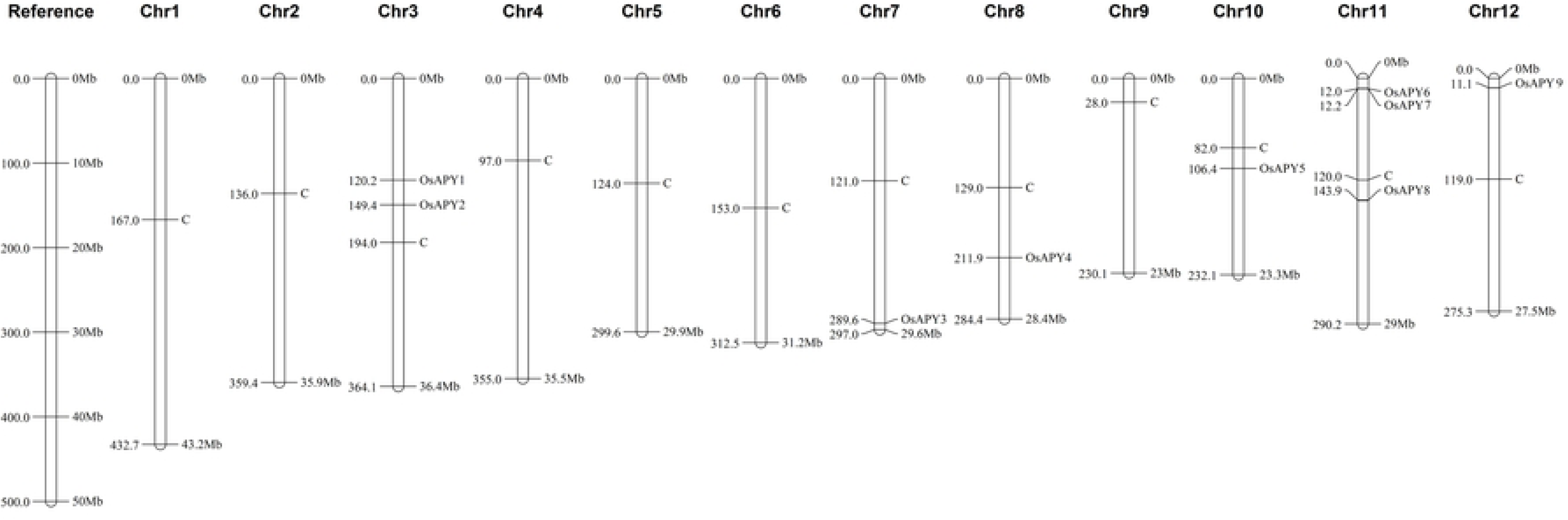
Distribution of *OsAPY*s throughout the rice genome. The figure was generated using MapChart. Chromosome number is presented on top. C depicts the centromere position. The position of every gene can be easily compared following the reference chromosome at the left.

### Identification of miRNAs targeting *OsAPY* genes

In this study, 103 potential, as well as unique miRNAs of 19–24 nucleotides in length, were found to target the *OsAPY* family members in rice. The highest number of miRNAs targeted *OsAPY9*, and *OsAPY3* was targeted by the lowest number. Fig 6 illustrates the regulatory relationships involving potential miRNAs as well as their targeted *APY*s. The majority of the miRNAs expected to target *OsAPY*s have a strong inhibitory effect through cleavage. The inhibitory action for only a few numbers of miRNAs was translation. The list of miRNAs targeting different *OsAPY*s and their mode of inhibition is given in the S4 Table.

**Fig 6.**
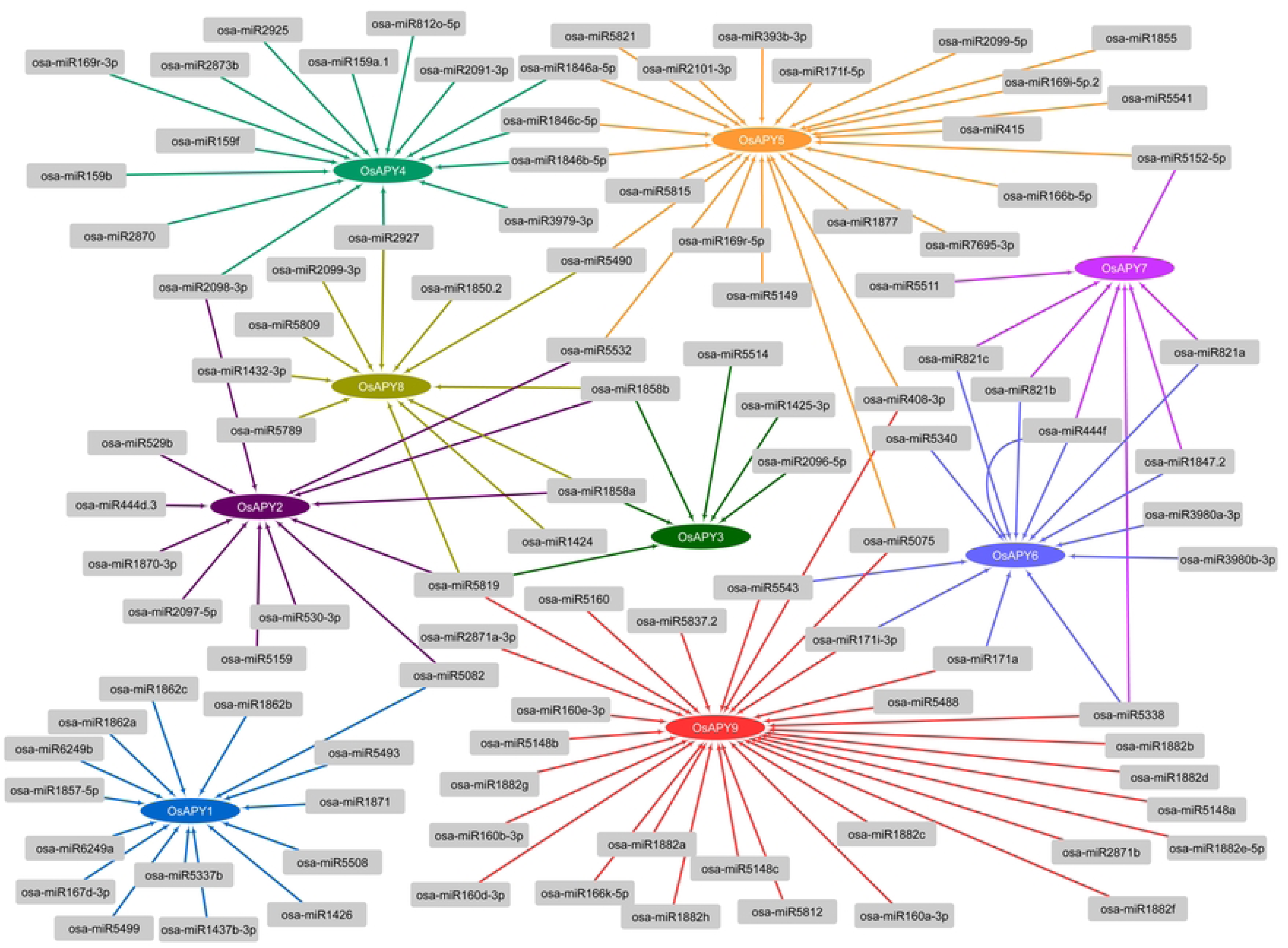
Identification of potential miRNAs targeting *OsAPY* genes. Visual depiction of miRNA-*OsAPY* interaction is generated via Cytoscape. Arrows depict the regulatory relationship, elliptical shapes of different colors represent the *OsAPY*s, and the miRNAs are shown in ash-colored boxes.

Among all the miRNAs, 23 miRNAs targeted more than one *OsAPY*s, and the rest were unique for each gene. It was found that most of the miRNAs had lower expression patterns. A heat map was generated (Fig 7) by comparing miRNA expression levels from PmiRExAt [73] across tissues and under a number of different abiotic stress conditions. Among all the miRNAs, osa-miR159a.1 targeting *OsAPY4* was expressed highly in all stresses and tissues except anther and leaves during the flowering stage. The target of osa-miR5532 was *OsAPY2* and *OsAPY5*, which was abundantly expressed in anther. osa-miR3979-3p, which targeted *OsAPY4*, was strongly expressed in roots. osa-miR408-3p targeting *OsAPY5*, *9*, was highly expressed in leaves during the flowering stage and in seedlings exposed to H_2_O_2_. These expressions of most of the miRNAs were downregulated during different stresses suggesting the increased expression of the *OsAPY*s in such stress conditions. Therefore, sequence-specific miRNA-mediated interaction may have a vital function in regulating *OsAPY* genes, which in turn may help plants act on environmental as well as growth signals.

**Fig 7.**
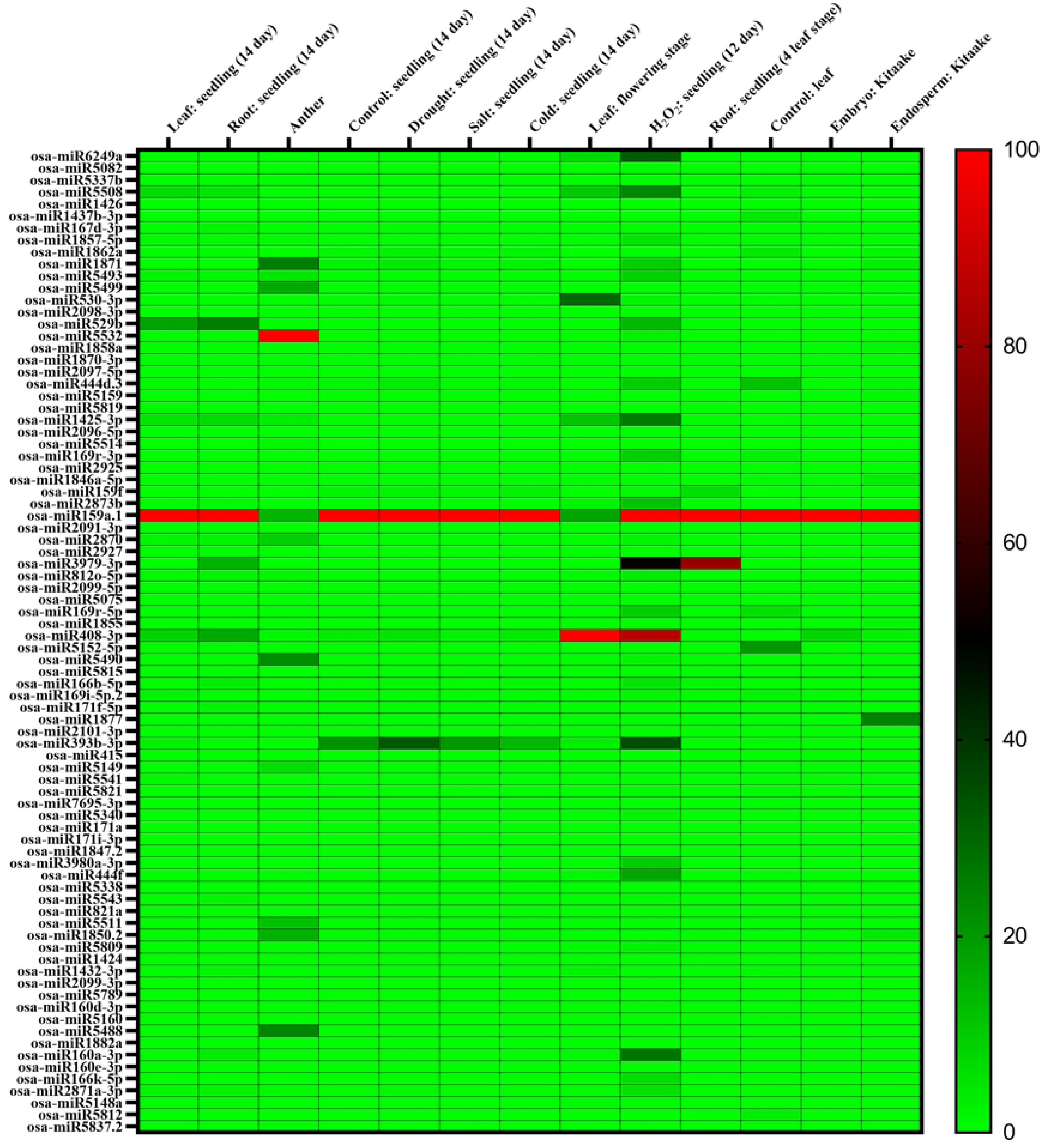
Analysis of *APY*-targeting miRNA expression in different abiotic stresses and tissues. The expression profile of each miRNA can be analyzed following the scale at the right. A heat map was generated using GraphPad Prism 9.0.0.

### Study of SNPs in *OsAPY*s

Rice SNP-Seek database [76] was used to better understand the variations in alleles of *OsAPY* members throughout 11 rice varieties and were selected based on their stress responses. It allowed identifying the Single Amino acid Polymorphisms (SAPs) (Table 4). SAPs were identified in 6 *OsAPY*s, but not in *OsAPY1*, *5*, and *8*. It indicates that the genes of *OsAPY1*, *5*, and *8* are substantially conserved among rice genotypes. The indica varieties include NERICA- L-27, Pokkali, Rasi, Vandana, Swarna, and IR-64, while Azucena and NERICA-1 are tropical japonica varieties. GIZA 159, Nagina 22, and Pusa (Basmati 1) are the members of the temperate japonica, aus, and unassigned variety, respectively. The study revealed that indica rice varieties have a higher rate of SAPs than other rice varieties. 5 SAPs were identified in *OsAPY2*; *OsAPY3*, *7*, and *9* showed 3 SAPs, while *OsAPY4* had only one SAP, but *OsAPY6* contained 4 SAPs.

**Table 4.** Distribution of single amino acid polymorphisms (SAPs) throughout the *APY*s of chosen rice varieties.

| Gene Name | Nipponbare | Pokkali | Rasi | IR-64 | NERICA-L-27 | Pusa (Basma ti 1) | Swarna | Vandana | NERICA-1 | Nagina 22 | Azucena | GIZA 159 |
| --- | --- | --- | --- | --- | --- | --- | --- | --- | --- | --- | --- | --- |
| *OsAPY2* | A<sub>109</sub> | A<sub>109</sub> | A<sub>109</sub> | A<sub>109</sub> | A<sub>109</sub> | A<sub>109</sub> | V<sub>109</sub> | A<sub>109</sub> | A<sub>109</sub> | A<sub>109</sub> | A<sub>109</sub> | A<sub>109</sub> |
|  | R<sub>258</sub> | R<sub>258</sub> | R<sub>258</sub> | R<sub>258</sub> | R<sub>258</sub> | R<sub>258</sub> | R<sub>258</sub> | R<sub>258</sub> | R<sub>258</sub> | R<sub>258</sub> | R<sub>258</sub> | H<sub>258</sub> |
|  | K<sub>295</sub> | K<sub>295</sub> | K<sub>295</sub> | K<sub>295</sub> | K<sub>295</sub> | K<sub>295</sub> | K<sub>295</sub> | K<sub>295</sub> | K<sub>295</sub> | N<sub>295</sub> | K<sub>295</sub> | K<sub>295</sub> |
|  | S<sub>365</sub> | A<sub>365</sub> | A<sub>365</sub> | S<sub>365</sub> | S<sub>365</sub> | S<sub>365</sub> | A<sub>365</sub> | A<sub>365</sub> | A<sub>365</sub> | S<sub>365</sub> | S<sub>365</sub> | S<sub>365</sub> |
|  | K<sub>388</sub> | Q<sub>388</sub> | Q<sub>388</sub> | Q<sub>388</sub> | Q<sub>388</sub> | Q<sub>388</sub> | Q<sub>388</sub> | Q<sub>388</sub> | Q<sub>388</sub> | Q<sub>388</sub> | K<sub>388</sub> | K<sub>388</sub> |
| *OsAPY3* | E<sub>111</sub> | A<sub>111</sub> | A<sub>111</sub> | A<sub>111</sub> | A<sub>111</sub> | A<sub>111</sub> | A<sub>111</sub> | A<sub>111</sub> | E<sub>111</sub> | A<sub>111</sub> | - | E<sub>111</sub> |
|  | N<sub>114</sub> | K<sub>114</sub> | K<sub>114</sub> | K<sub>114</sub> | K<sub>114</sub> | K<sub>114</sub> | K<sub>114</sub> | K<sub>114</sub> | N<sub>114</sub> | K<sub>114</sub> | N<sub>114</sub> | N<sub>114</sub> |
|  | V<sub>136</sub> | A<sub>136</sub> | A<sub>136</sub> | A<sub>136</sub> | A<sub>136</sub> | A<sub>136</sub> | A<sub>136</sub> | A<sub>136</sub> | V<sub>136</sub> | A<sub>136</sub> | V<sub>136</sub> | V<sub>136</sub> |
| *OsAPY4* | S<sub>207</sub> | G<sub>207</sub> | G<sub>207</sub> | G<sub>207</sub> | G<sub>207</sub> | G<sub>207</sub> | G<sub>207</sub> | G<sub>207</sub> | G<sub>207</sub><br>S<sub>207</sub> | G<sub>207</sub> | G<sub>207</sub> | S<sub>207</sub> |
| *OsAPY6* | M<sub>77</sub> | K<sub>77</sub> | K<sub>77</sub> | K<sub>77</sub> | K<sub>77</sub> | K<sub>77</sub> | K<sub>77</sub> | K<sub>77</sub> | M<sub>77</sub> | - | M<sub>77</sub> | M<sub>77</sub> |
|  | I<sub>162</sub> | R<sub>162</sub> | R<sub>162</sub> | R<sub>162</sub> | R<sub>162</sub> | R<sub>162</sub> | R<sub>162</sub> | R<sub>162</sub> | I<sub>162</sub> | R<sub>162</sub> | I<sub>162</sub> | I<sub>162</sub> |
|  | P<sub>252</sub> | S<sub>252</sub> | S<sub>252</sub> | S<sub>252</sub> | - | S<sub>252</sub> | S<sub>252</sub> | S<sub>252</sub> | P<sub>252</sub> | S<sub>252</sub> | P<sub>252</sub> | P<sub>252</sub> |
|  | E<sub>261</sub> | K<sub>261</sub> | K<sub>261</sub> | K<sub>261</sub> | K<sub>261</sub> | K<sub>261</sub> | K<sub>261</sub> | K<sub>261</sub> | E<sub>261</sub> | K<sub>261</sub> | E<sub>261</sub> | E<sub>261</sub> |
| *OsAPY7* | I<sub>236</sub> | T<sub>236</sub> | T<sub>236</sub> | T<sub>236</sub> | T<sub>236</sub> | T<sub>236</sub> | T<sub>236</sub> | T<sub>236</sub> | I<sub>236</sub> | T<sub>236</sub> | I<sub>236</sub> | I<sub>236</sub> |
|  | E<sub>242</sub> | G<sub>242</sub> | G<sub>242</sub> | G<sub>242</sub> | G<sub>242</sub> | G<sub>242</sub> | G<sub>242</sub> | G<sub>242</sub> | E<sub>242</sub> | G<sub>242</sub> | E<sub>242</sub> | E<sub>242</sub> |
|  | V<sub>264</sub> | V<sub>264</sub> | V<sub>264</sub> | V<sub>264</sub> | V<sub>264</sub> | V<sub>264</sub> | V<sub>264</sub> | V<sub>264</sub> | I<sub>264</sub> | V<sub>264</sub> | V<sub>264</sub> | V<sub>264</sub> |
| *OsAPY9* | N<sub>75</sub> | N<sub>75</sub> | D<sub>75</sub> | N<sub>75</sub> | N<sub>75</sub> | N<sub>75</sub> | N<sub>75</sub> | N<sub>75</sub> | N<sub>75</sub> | D<sub>75</sub> | N<sub>75</sub> | N<sub>75</sub> |
|  | Q<sub>290</sub> | H<sub>290</sub> | Q<sub>290</sub> | H<sub>290</sub> | H<sub>290</sub> | H<sub>290</sub> | H<sub>290</sub> | H<sub>290</sub> | H<sub>290</sub> | Q<sub>290</sub> | H<sub>290</sub> | Q<sub>290</sub> |
|  | N<sub>350</sub> | N<sub>350</sub> | D<sub>350</sub> | N<sub>350</sub> | N<sub>350</sub> | N<sub>350</sub> | N<sub>350</sub> | N<sub>350</sub> | N<sub>350</sub> | D<sub>350</sub> | N<sub>350</sub> | N<sub>350</sub> |

- Represent the polymorphism in the non-coding region.

### Analysis of secondary and tertiary structure

The secondary structure analysis revealed the position of alpha-helix, beta-sheet, isolated beta bridge, turn, coil, 310-helix, and transmembrane helix (Fig 8). Their percentage and the position of the transmembrane helix are given in the S5 Table. It revealed that alpha-helix is dominant over all the other secondary structures, followed by the turn, beta-sheet, coil, and 310-helix. There were some exceptions in OsAPY2, 4, 5, and 8. In OsAPY5, the coil percentage is higher than the beta-sheet, and in OsAPY2, it is higher than the turn and beta- sheet. In OsAPY4 and OsAPY8, the beta-sheet percentage is higher than that of turn. The higher percentage of helix and beta-sheet indicate the stability of the OsAPYs [132], and the structures like random coils are critical for the signaling cascades [133]. The analysis of the motifs spanning the membrane (MSM) indicated that 4 OsAPYs contained one membrane- spanning motif (MSM), 4 OsAPYs contained 2 MSMs, and 1 OsAPY contained no MSM. OsAPY1, 3, 7, 8 contained only one membrane-spanning motif at N-terminal. OsAPY2, 4, 5, and 6 had two MSMs positioned at the N- and C-terminals separately except OsAPY6, where both the MSMs were located on N-terminal. On the other hand, OsAPY9 contained no MSM.

**Fig 8.**
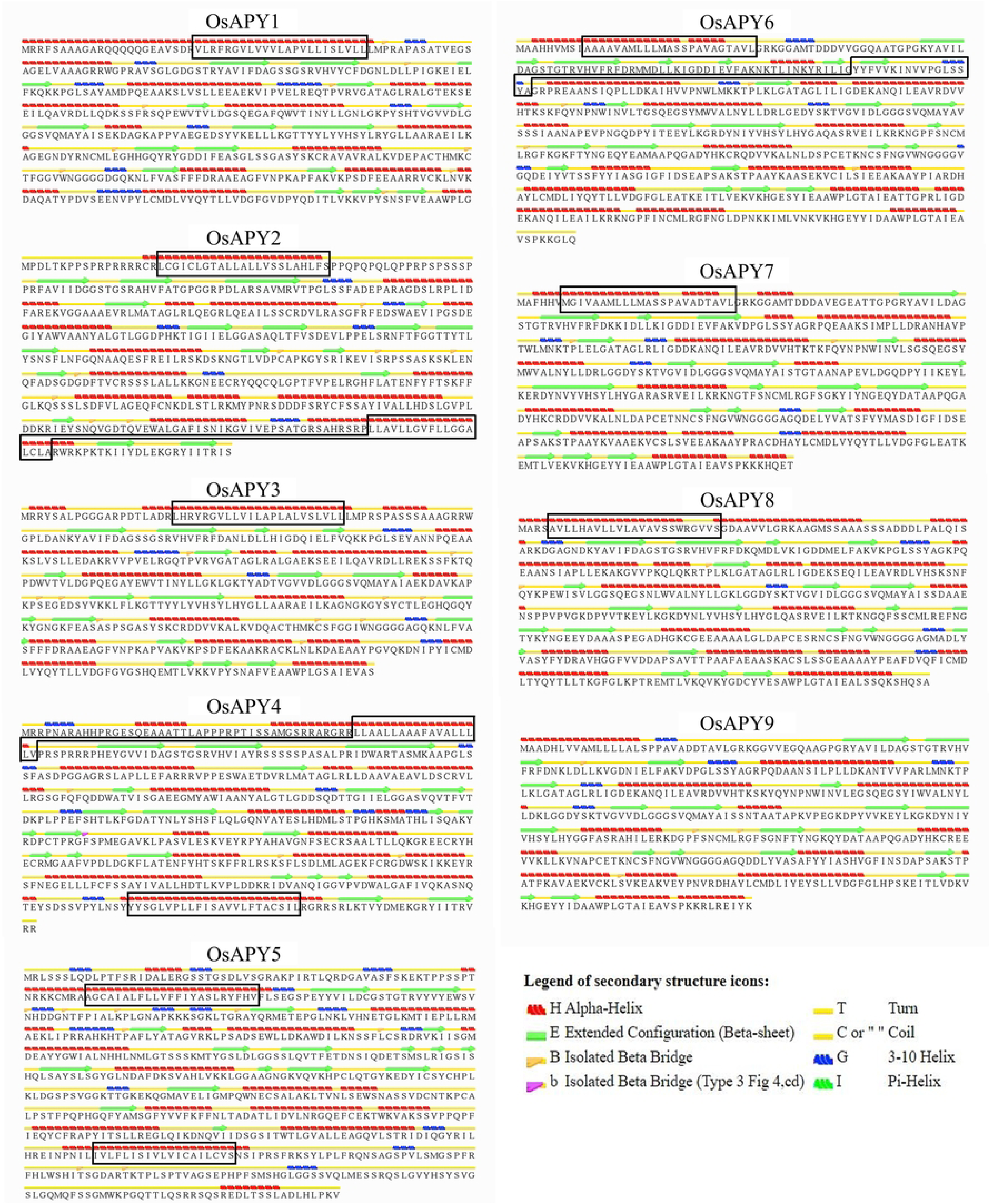
Secondary structure analysis of the nine rice APYs. Secondary structures were generated using the STRIDE program, and the corresponding positions of membrane-spanning motifs were identified via the TMHMM server. The cross-membrane domains were marked with a black colored box. Legend at the bottom-right side, shows the icon foe each secondary structure.

RoseTTAFold, a deep learning-based modeling approach, was applied to construct 3D models of proteins using the Robetta web server [79]. The generated models were further refined by GalaxyRefine [80], and then the energy minimization was done via Swiss-PdbViewer [81]. Fig 9 illustrates the modeled tertiary structures of all the OsAPY proteins, visualized via Discovery Studio. These generated structures were then subjected to validation analysis utilizing PROCHECK [82], ERRAT [83], and ProSA-web server [84] (S6 Table). Ramachandran plot analysis in PROCHECK was used to evaluate protein quality [82]. In the favored and additional allowed regions, over 90% of the residues were located, with only <1.5% in the disallowed regions, which confirmed that the projected models are of good quality (S2 Fig). The proteins exhibited an overall quality factor of >87, according to ERRAT [83] analysis (S3 Fig). ProSA-web [84] revealed the Z-score, the level of nativeness of the designed models (S6 Table), and the energy plot, which shows the local quality of the models. The Z-score was found to be in the spectrum of values generally reported for native proteins, implying higher quality of the structures generated, and the models were well within the range of X-ray crystal structure (S4 Fig). As per the energy plot (S5 Fig), all the residues in the simulated structure had a lower value of energy. These findings indicate the excellent quality of the tertiary structures of modeled proteins.

**Fig 9.**
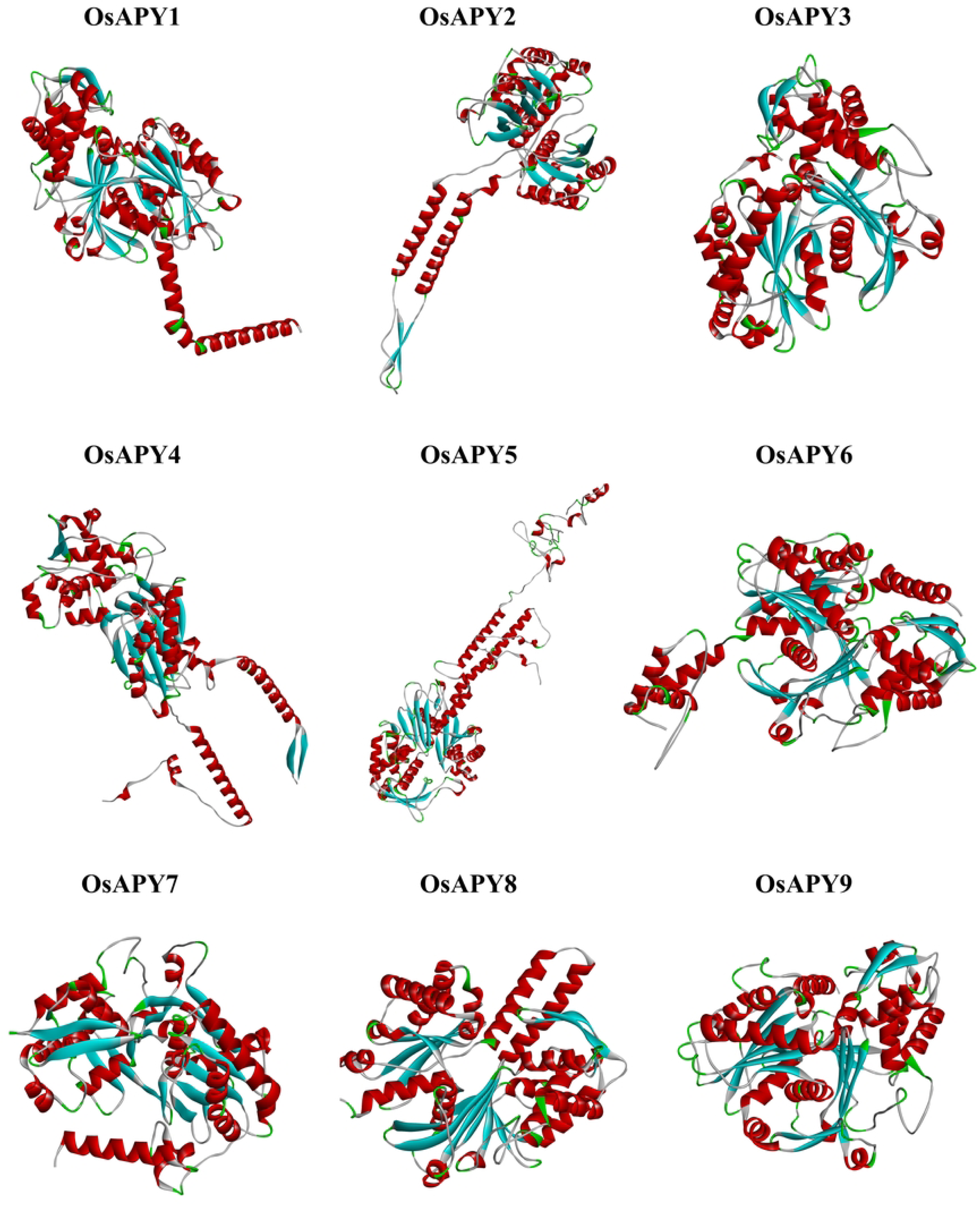
Tertiary structure analysis of the nine rice APYs. The online tool Robetta was used to construct the tertiary structure, and these structures were visualized via Biovia Discovery Studio Visualizer.

### Analysis of molecular docking

ATP, a vital ligand of APY proteins, was chosen for the analysis of docked protein-ligand complex. The docking analysis was done using the HDOCK server, and it revealed that the docking score of the protein and ATP was between -199.05 and -152.53 (S7 Table). This optimal docking indicated an excellent binding affinity. According to PDBSum [87] analysis, the protein and ATP complex contained 3 to 11 hydrogen bonds (S8 Table). The interaction between the protein and ATP visualized via Discovery Studio [85] revealed the presence of different types of H-bond, electrostatic bond, and hydrophobic bonds (Fig 10 and S9 Table). Validation of the protein-ligand complex via PROCHECK and ERRAT suggested the high quality of the complexes (S6 and S7 Figs).

**Fig 10.**
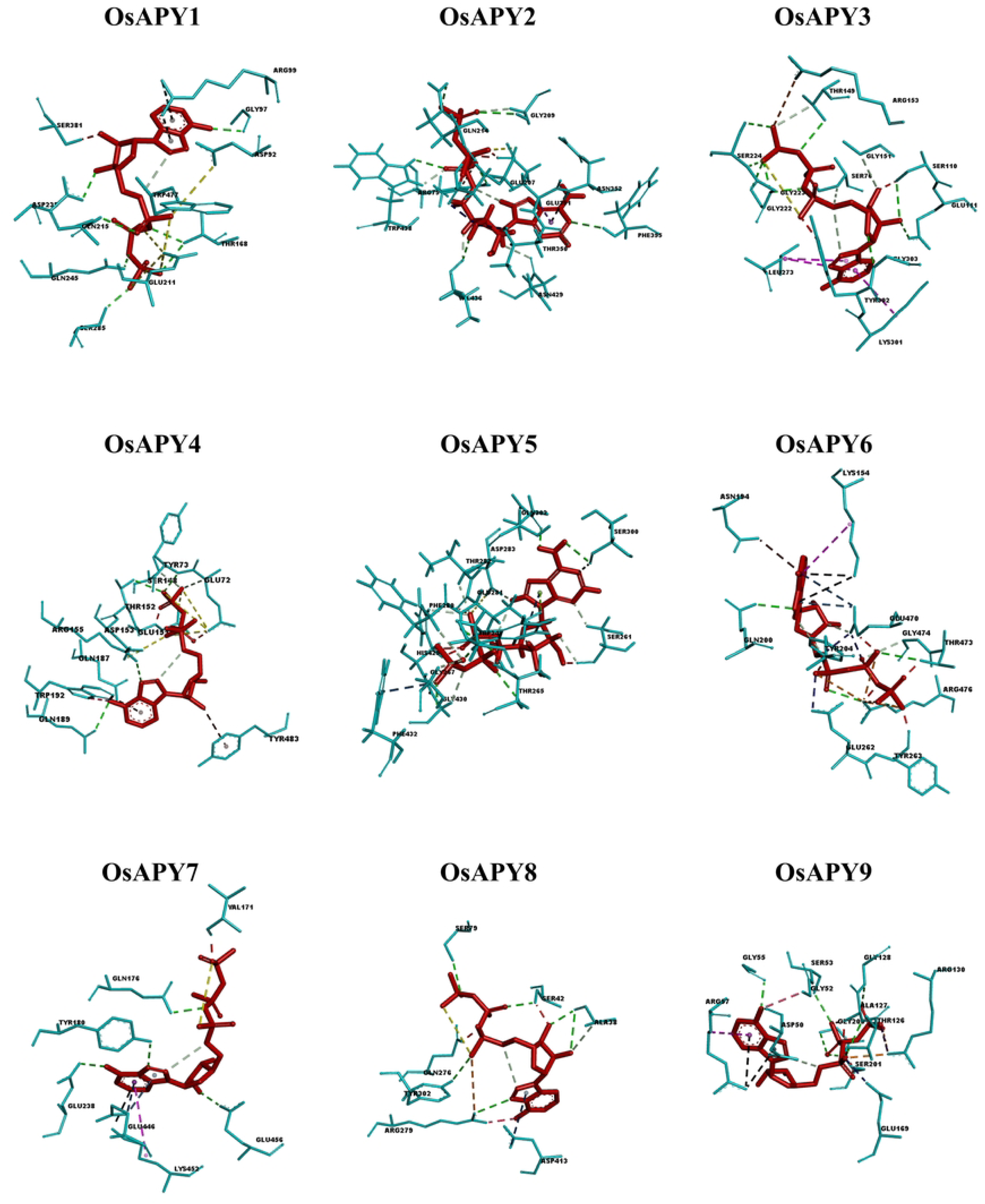
Protein-ligand residual interaction analysis of OsAPYs with ATP depicted by aqua and red color, respectively. Illustration of bonds: carbon-hydrogen bonds- snowy mint lines, conventional hydrogen bonds- green lines, pi-cation bonds- black lines, unfavorable bumps- red lines, unfavorable negative-negative interactions- yellow lines, attractive charge interactions- orange lines, pi-donor hydrogen bonds- deep brown lines, pi-sigma bonds- light purple lines, unfavorable acceptor- acceptor interactions- navy blue lines, pi-alkyl bonds- magenta lines, unfavorable donor-donor interactions- pink lines, pi-anion bonds- deep-sea blue lines, pi-lone pair interactions- lemon green lines, salt bridge- purple lines.

OsAPY1 formed seven conventional hydrogen bonds, 2 Pi-cation interactions, and seven unfavorable interactions (Fig 10). Among all the protein residues, THR168 formed 2 of the seven hydrogen bonds, and ARG99 formed both the Pi-cation interactions (S9 Table). OsAPY2 had one attractive charge interaction, 1 Pi-donor hydrogen bond, six conventional hydrogen bonds, five unfavorable interactions, six carbon-hydrogen bonds, and 1 Pi-sigma interaction with ATP. OsAPY3 made one attractive charge interaction, three Pi-alkyl bonds, carbon-hydrogen bonds, two unfavorable interactions, and nine conventional hydrogen bonds. OsAPY4 created four conventional hydrogen bonds, 2 Pi-donor hydrogen bonds, and two carbon-hydrogen bonds. OsAPY5 formed ten carbon-hydrogen bonds, six conventional hydrogen bonds, 1 Pi-lone pair interaction, 2 Pi-anion interactions, and seven unfavorable interactions. OsAPY6 made 2 Pi-cation interactions, Pi-donor hydrogen bonds, carbon-hydrogen bonds, Pi-anion interactions, four conventional hydrogen bonds, four attractive charge interactions, and 1 Pi-alkyl interaction. OsAPY7 and OsAPY8 formed 9 and 12 interactions with ATP, respectively. OsAPY9 had one salt bridge interaction, two attractive charge interactions, seven conventional hydrogen bonds, 2 Pi-cation interactions, 1 Pi-anion interaction, 1 Pi-alkyl interaction, and eight unfavorable interactions. These bonds between the OsAPYs and ATP indicate a strong docking interaction.

### Investigation of the expression profiles of *OsAPY* genes using RNA-seq data

Using RNA transcript profiling to analyze the gene expression is an efficient technique. To better comprehend the expression profile of *OsAPY*s, the RNA-seq data was utilized under several tissue types and stresses (Fig 11). The Rice Genome Annotation Project [52] provided RNA-seq data for *OsAPY* expression profiling in several tissues which was used to produce a heatmap (Fig 11).

**Fig 11.**
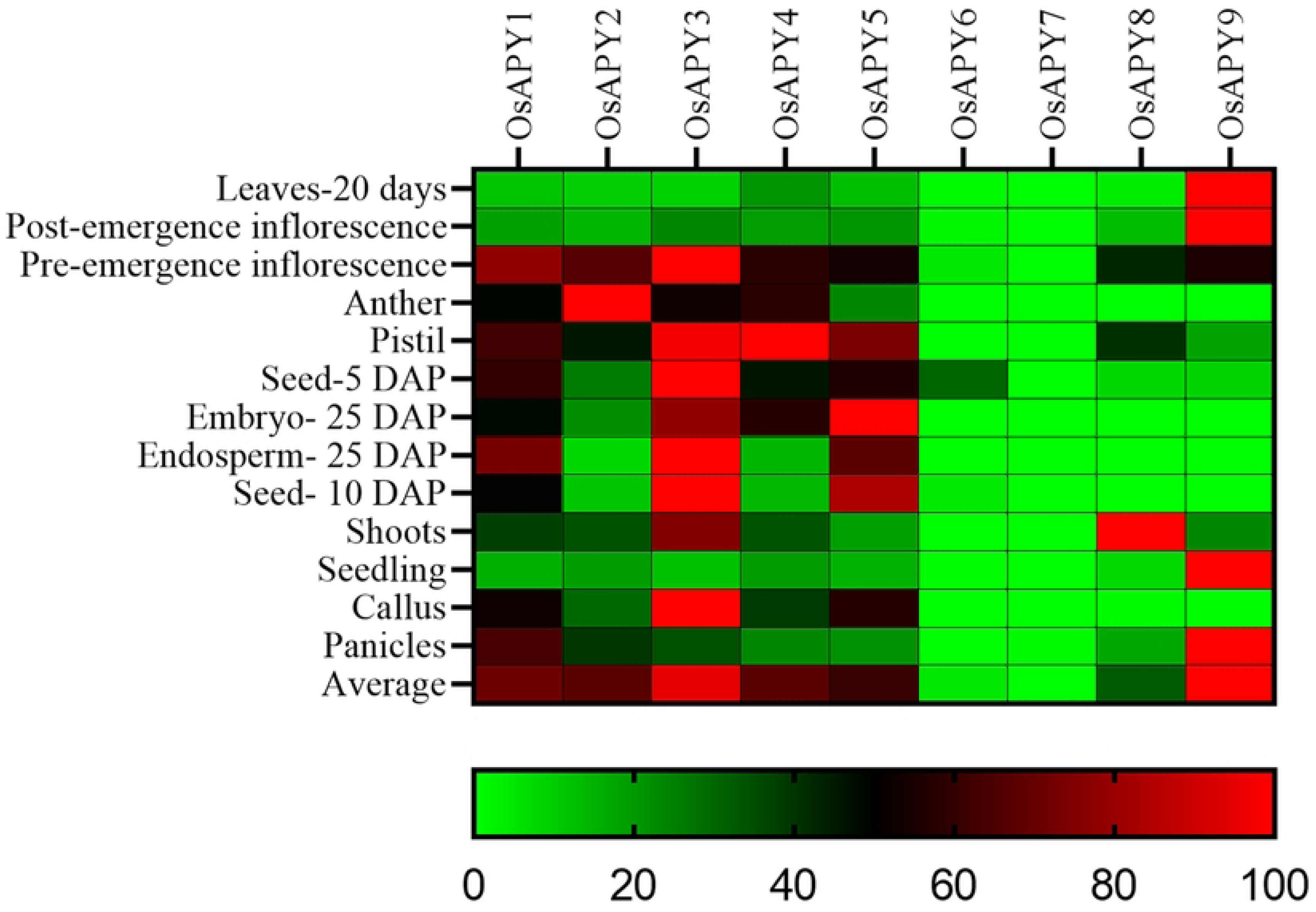
Heatmap of the expression pattern of the *OsAPY* genes according to tissue type. The RNA-seq data collected from the Rice Genome Annotation Project was used to conduct expression analysis. The heat map was generated using GraphPad Prism 9.0.0. Expression profile can be analyzed following the bottom located scale. The red boxes represent high expression rates, greens depict low expression rates, and black boxes signify moderate expression levels.

The analysis revealed that the expression was higher in the inflorescence, seed, pistil, embryo, and endosperm. *OsAPY1* and *3* upregulated in most of these tissues, whereas in the pre-emergence inflorescence and anther, *OsAPY2* and *4* upregulated. *OsAPY4* was expressed highly in the pistil. For *OsAPY5*, it was higher in pistil, seed, embryo, and endosperm among these tissues. Some of these showed an expression pattern that was tissue-specific, including *OsAPY2*, upregulated in anther, inflorescence, and downregulated in the other types of tissues. In the same manner, the expression of *OsAPY8* was high in shoots. All the *OsAPY*s were downregulated in leaves, post-emergence inflorescence, and seedlings except *OsAPY9*, which was upregulated in these tissues. In shoot tissue, *OsAPY*3 and *8* were highly expressed; on the other hand, *OsAPY3* and *5* were upregulated in callus, and it was *OsAPY1* and *9* in panicles.

The profile of expression from RNA-seq data of *OsAPY*s was investigated to reveal their role in response to different stresses. Their expression profile revealed expression in abiotic and biotic stresses. Figs 12 and 13 show a heat map of their expression profile in biotic and abiotic stress. There was no pre-analyzed RNA-seq data of *OsAPY9*, so the gene expression data of only eight genes were analyzed.

**Fig 12.**
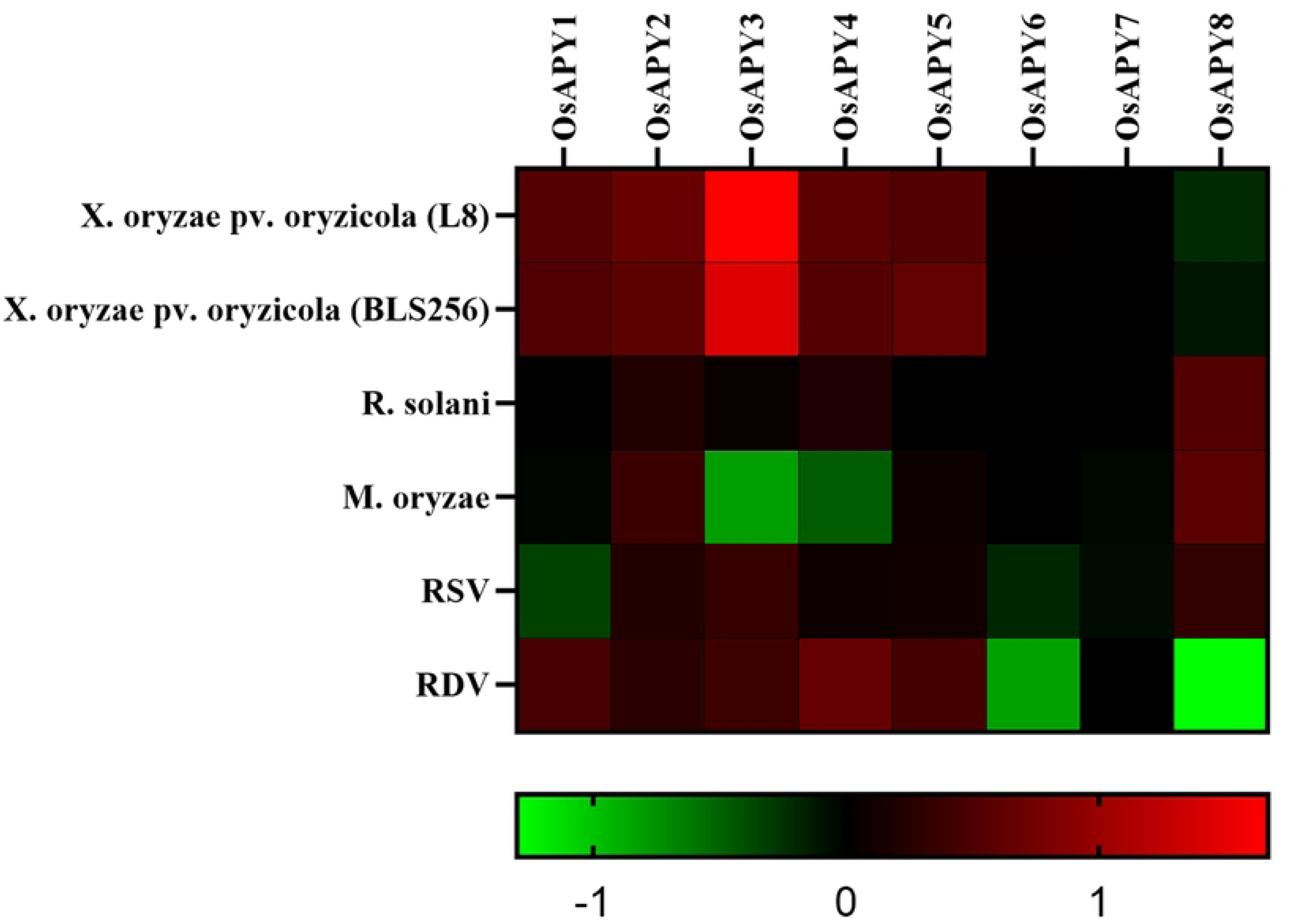
Heatmap of *OsAPY* expression profile in response to various biotic stress conditions. Expression analysis was done utilizing the RNA-seq values of relative expression from GENEVESTIGATOR and using GraphPad Prism 9.0.0, the heat map was generated. The expression profile could be analyzed according to the scale just at the base. The red boxes represent upregulated expression; greens depict downregulated expression, and black boxes signify no change in expression levels.

**Fig 13.**
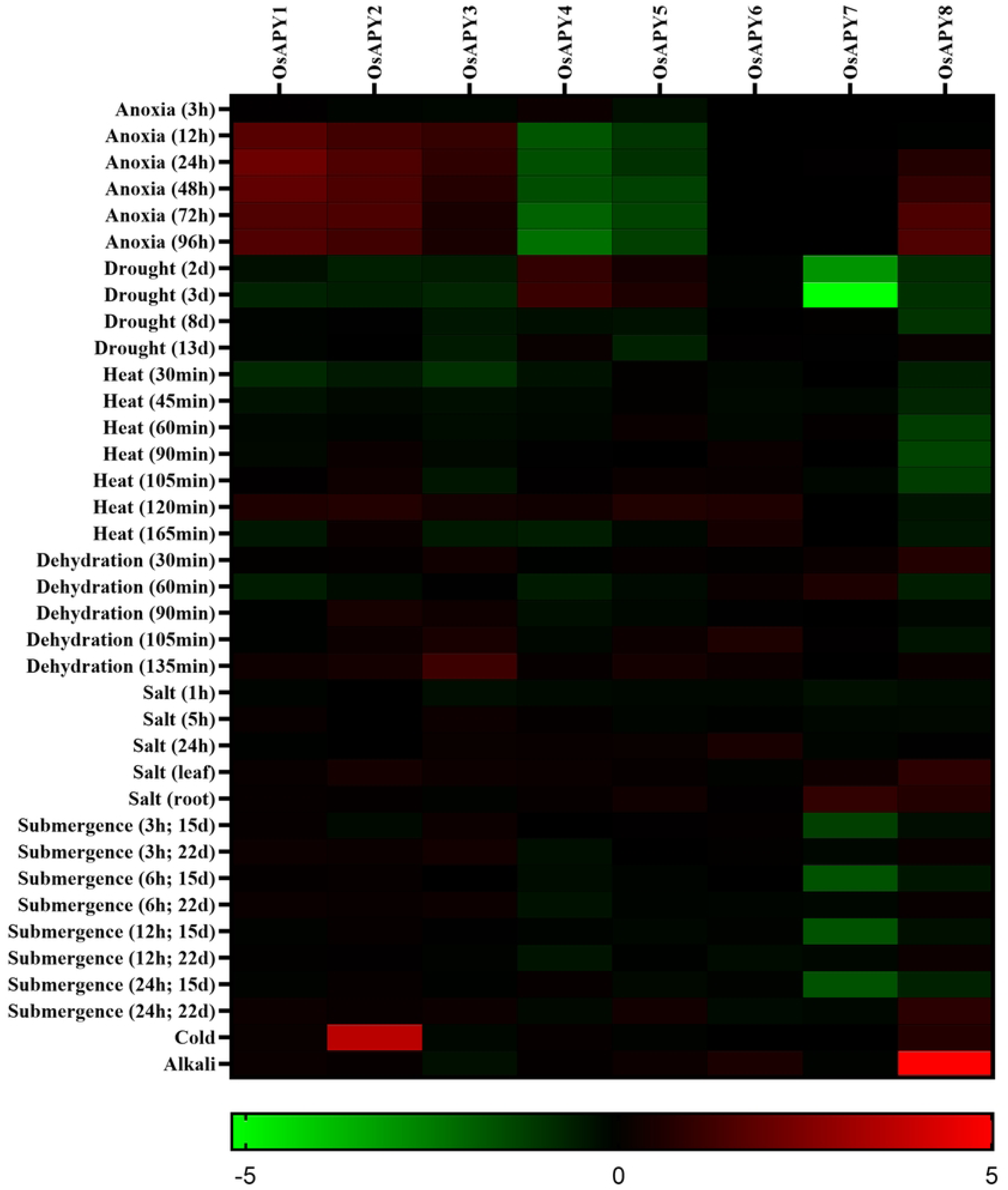
Heatmap of *OsAPY* expression profile in response to various abiotic stress conditions. Expression analysis was done using the RNA-seq values of relative expression from GENEVESTIGATOR. The heat map was generated using GraphPad Prism 9.0.0. The expression profile could be analyzed according to the scale just at the base. The red boxes represent upregulated expression; greens depict downregulated expression, and black boxes signify no change in expression levels.

In response to infection by two strains of rice bacterial leaf streak pathogen (*Xanthomonas oryzae* pv. *oryzicola)*, *OsAPY1-5* were upregulated, but *OsAPY8* was downregulated (Fig 12). This finding addressed the role of *OsAPY*s in managing rice stress responses against bacterial leaf streak pathogens. It was observed that the infection with *Rhizoctonia solani* upregulated the expression of *OsAPY2*, *4*, and *8*. Rice blast fungus (*Magnaporthe oryzae*) inoculation induced the expression of *OsAPY2*, *5*, and *8* indicating their possible role in rice blast disease. Infection with rice dwarf virus (RDV) upregulated the expression of *OsAPY1-5*; on the other hand, *OsAPY6*, *8* were downregulated. *OsAPY2*, *3*, *5*, and *8* got upregulated in response to rice stripe virus (RSV) infection, but at the same time, *OsAPY1* and *6* got downregulated.

During anoxic conditions, the expression of *OsAPY1-3* and *8* was upregulated, but *OsAPY4* and *5* were downregulated (Fig 13). *OsAPY4* showed a slight increase in expression only in 3 hours-long anoxic conditions. In *OsAPY8*, the level of expression was proportional to the duration of the anoxic condition. Rice subjected to drought for 2 and 3 days showed upregulation of *OsAPY4* and *5*, but all the other *OsAPY*s, especially *OsAPY7*, were downregulated except *OsAPY6*, which showed no change in expression. The extended period of drought condition downregulated the expression level. The gene expression level increased with time duration in heat stress, but after a specific time, it started to decrease. In the case of heat treatment for 120 minutes, the expression of *OsAPY1-6* was upregulated, but gene expression levels reduced during an extended period of heat treatment (165 minutes). This result indicated that *OsAPY*s displayed late-term responses to heat stress, but extended stress interrupted their expression. When subjected to dehydration, similar late-term expression was observed. 135 minutes long dehydration stress induced the expression of all the *OsAPY*s except *OsAPY4* and *7*. Under salt stress, the expression was relatively higher in leaves than in roots.

All the genes except *OsAPY1*, *5*, and *6* were upregulated in leaves, whereas in root tissues, only *OsAPY5*, *7*, and *8* were upregulated. Under submergence stress, the expression level got increased with the increase in time. As the heatmap depicted (Fig 13), after 22 days of submergence for 24 hours, there was an increase in the expression of *OsAPY1*, *3*, *5*, and *8*. *OsAPY5* and *8* had an increased expression after prolonged exposure to submerged conditions as there was no significant expression in the case of short-term exposure to submerged conditions. Alkali treatment upregulated the expression of *OsAPY5*, *6*, and especially of *8*, whereas for *OsAPY2* and *8*, the expression was induced in cold.

### Investigation of expression profile using RT-qPCR data in response to several abiotic stresses

Relative expression ratio of *OsAPY* genes in the rice seedling leaves was assessed under cold, cadmium, salinity, submergence, drought, and heat stress. Heat map of their real-time expression data in these different stress conditions after 18-20 hours is depicted in Fig 14.

**Fig 14.**
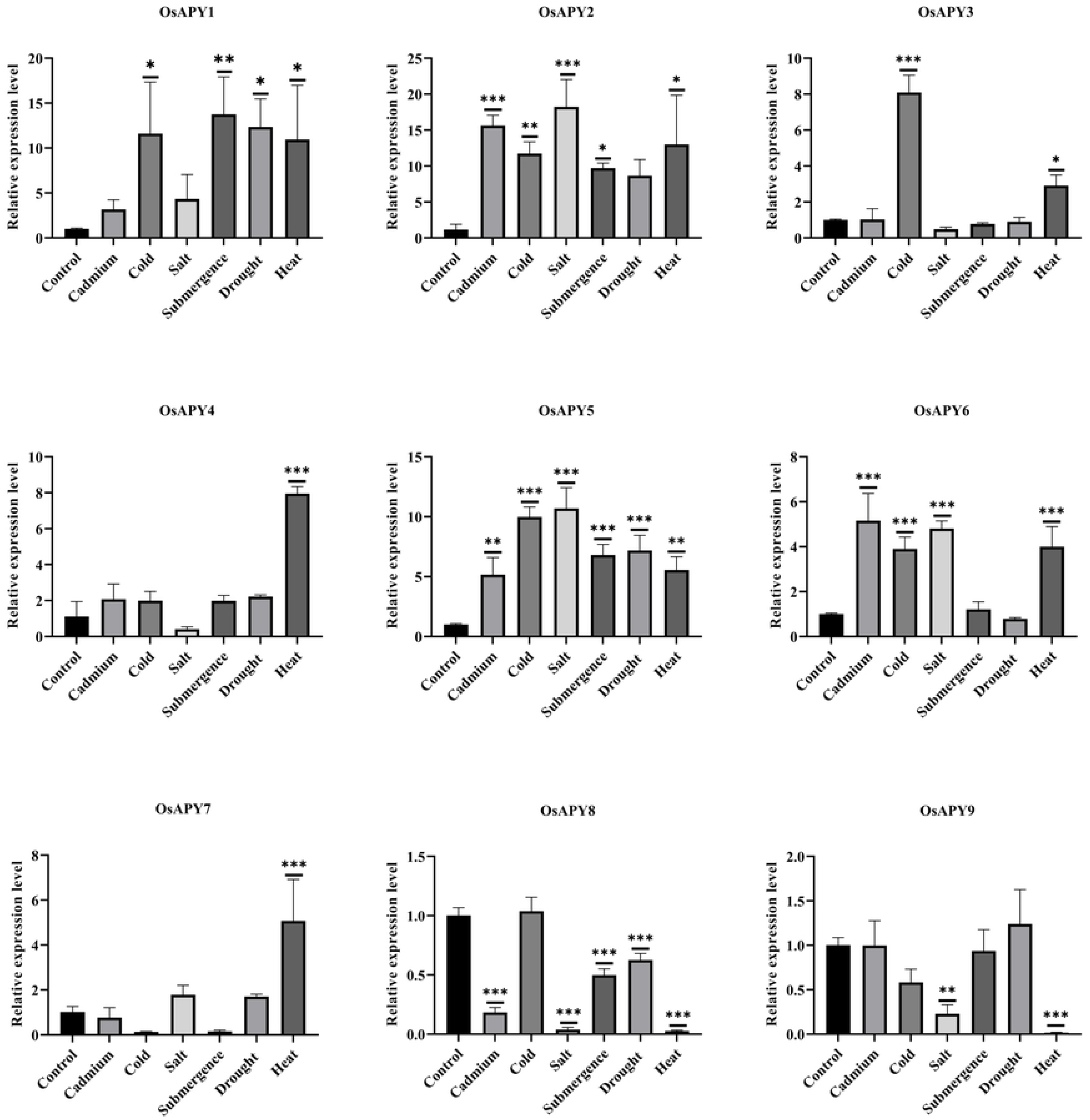
Analysis of relative expression of *OsAPY*s under various abiotic stresses. The Y-axis depicts the relative expression of each gene analyzed via RT-qPCR in the rice plant leaves. The X-axis indicates the various abiotic stress conditions under which the relative expression analysis was done. Technical replication was used to find the mean value of expression at different treatments. MS Office 365 and GraphPad Prism 9.0.0 were used to analyze the data. A one-way ANOVA, followed by a Bonferroni post hoc test, was used to assess the significant differences (P ≤ 0.05). To represent the significant differences, different means were labeled with different number of stars (*).

As shown in Fig 14, all the *OsAPY* genes except *OsAPY8* and *9* showed significantly higher expression under heat stress. Among all the stress environments, *OsAPY5* had considerably higher expression levels (more than five-fold) in all the stress conditions. The RNA-seq data (Fig 13) also demonstrated its enhanced expression during drought, submergence, dehydration, alkali, salt, and heat stress.

Cold, submergence, drought, and heat stress, all resulted in considerably more significant levels of *OsAPY1* expression. The expression level during submergence stress (more than ten-fold) was significantly higher, and the result obtained from RNA-seq data (Fig 13) corroborated this observation. Under cadmium, cold, salinity, submergence, and heat stress, *OsAPY2* demonstrated significant expression, and RNA-seq data revealed consistent outcomes (Fig 13), especially in cold conditions, its expression was very high. However, the real-time expression level was much more significant in response to cadmium and salinity stress. *OsAPY3* expression was substantially increased in heat and cold stress. Compared to the control condition, it was much greater during cold stress and demonstrated an 8-fold upregulation. Only heat stress raised the expression of *OsAPY4* (approximately eight-fold) and *7* (more than four-fold), whereas salinity and heat stress significantly downregulated the expression of *OsAPY9*. Under heat stress, RNA-seq data also revealed a higher level of *OsAPY4* expression. During cadmium, cold, salinity, and heat stress, *OsAPY6* expression was significantly enhanced, and RNA-seq data also exhibited upregulated expression under salinity and heat stress (Fig 13). Under cadmium, salinity, submergence, drought, and heat stress, *OsAPY8* demonstrated significant downregulation. Though the expression under cold conditions was higher, it was not significant. Its downregulated expression level during drought, heat, and submergence stress and the increased one in cold was also supported by RNA-seq data. According to RNA-seq data, prolonged exposure to submerged situations increased its expression. This result indicates a potential function for *OsAPY*s in regulating abiotic stress.

## Discussion

A significant role is played by apyrase (*APY*) in regulating the growth of plants, developmental changes, and different stresses [18, 24]. When the plants encounter any stress, the extracellular ATP (eATP) level rises which in turn increases ROS expression, triggers several other stress-induced genes, and can also cause apoptosis [5, 17]. Apyrase can hydrolyze eATP [134], and thus it has a major function in regulating eATP-mediated ROS production, and cellular apoptosis and in providing tolerance towards various stress conditions. The majority of the world takes rice as their main diet, and Asia alone contributes 90% to global rice production [135]. Different stresses are blamed for yield losses in commercially essential crops in many parts of the world. So, manipulating *APY*s can be a great step towards developing genetically modified multiple stress-tolerant rice and improving its annual production as *APY*s are associated with various developmental stages as well as stress responses of the plants. For this, the *APY* gene family should be studied intensively. Wheat and *Arabidopsis* have been extensively investigated for this gene family [2, 23], but an investigation in rice is still not conducted. This study was designed to identify, characterize, and for analyzing the patterns of *APY* expression in rice.

Rice was found to have nine apyrase genes, as per this study, and all of these genes contained all the 5 ACRs [22]. An expanded insight into the significance of apyrase under abiotic and biotic stress was gained by the identification, characterization, and profiling of the expression of *APY* genes. These genes are different in their sequence length, molecular weight, pI value, and GRAVY value which means that this gene family is diverse. The GRAVY value of the proteins is negative, confirming the proteins are hydrophilic [92].

Ecto-APYs are usually present on plasma membrane and other cell surfaces, and endo-APYs are localized on ER, Golgi body, etc. [23, 136], and extracellular ATP is regulated by both endo- and ecto-APYs [19, 137]. In this investigation, the proteins were present on the plasma membrane, mitochondria, endoplasmic reticulum, and chloroplast, suggesting that both endo-APYs and ecto-APYs are present in rice [2, 23]. Nevertheless, the localization of OsAPY3, 4, and 9 varies according to the two different tools. Findings of CELLO [58] indicated there are three ecto-APYs (OsAPY2, 4, 5), but WoLF PSORT [57] suggested there are two ecto-APYs (OsAPY2, 5). It will take further research to find their precise position.

During evolution, gene duplication has a significant contribution in the process of gene expansion., and during plant development and growth, gene duplication can assist plants in adapting to various conditions [138]. The study of duplication events identified six duplication occurrences within the genome. The evolution of these genes occurred under the influence of purifying selection, which was suggested by the value of Ka/Ks, which was less than 1 [93]. These duplications occurred in genes situated on different chromosomes, indicating the segmental type of duplication being the main force of diversification [93]. Although segmental duplication preserves the primary functional group, it leads to differentiation in the manner of duplicated genes [139].

*OsAPY*s were related to *APY*s from *Arabidopsis* and wheat by constructing a phylogenetic tree in order to better understand their structure and functions. It was found that the phylogenetic tree was composed of three distinct groupings. According to earlier research on *Arabidopsis* and wheat *APY*s, this conclusion is in accordance [2, 23]. None of the *OsAPY*s was categorized as members of any new group. *OsAPY* was found to be a very archaic family of genes, originating before even the splitting of monocotyledon (rice and wheat) and dicotyledon (*Arabidopsis*) plants because the *OsAPY* genes shared an equal number of groups with the *AtAPY* genes and with the *TaAPY* genes [140].

The number and orientation of exon-introns in plant genes play an essential role in evolution [141]. Gene structure analysis predicted that there was no intronless gene which indicates their high expression, and this gene family is an ancient one and not recently evolved [142, 143].

*OsAPY5* contained the lowest exon and intron, and *OsAPY6* contained maximum exons and introns. Although *OsAPY1*, *3*, *6* belonged to group I of the phylogenetic tree, their exon-intron structure was different from the other members of group I. *OsAPY2* and *OsAPY4* belonged to group II, but their gene structure was consistent with the other members of group I. *OsAPY5* which belonged to group III had a completely different exon-intron structure.

Conserved motifs are essential for proteins to be functional; and for their specificity. In order to better understand how proteins interact, it is helpful to find the common patterns in structure among them [144]. Conserved motif analysis revealed that motif 1, 5, 8, and 10 were conserved across all OsAPYs; motif 2 was present in all the OsAPYs except OsAPY2. Motif 3, 4, 6, 7, and 9 were the least distributed. These were present only in group I of phylogenetic tree, and the members of each phylogenetic group had an almost similar structure. Their function analysis revealed that 8 of the identified motifs are members of the GDA1-CD39 family, which confirms that the identified genes are members of this GDA1-CD39 nucleosidephosphatase superfamily.

The percentage of alpha-helix, beta-sheet, isolated beta bridge, turn, coil, and 310-helix was similar in all the proteins. The secondary structure analysis showed the dominance of alpha-helix over other structures, which implied the stability of the protein structure [132]. The membrane-spanning motif numbers varied from 0 to 2, which are similar to the previous findings [2, 23]. Proteins containing one motif had only N- terminal MSM. The proteins with two motifs had both N- and C- terminal MSMs except OsAPY6, which had only 2 N-terminal MSMs. N- as well as C- terminal MSMs are found in OsAPY2, 4, and 5. These proteins with both the N- and C- terminal MSMs were considered as ecto-APYs according to CELLO [58]. In the earlier studies on human APYs, proteins containing both the MSMs were also predicted to be ecto-APYs [2]. This finding is in accordance with the result obtained from CELLO [58] and TMHMM web servers [78].

The tertiary structure analysis revealed that the predicted structures of all the proteins were of good quality, as confirmed by various tools. These structures could be utilized in analyzing the proteins more precisely in future research. Their docking analysis showed that there existed four categories of bonds between the proteins and their ligand ATP. The high numbers of hydrogen bonds corroborate the stability of the protein-ligand complex, as the stability of such complexes is typically facilitated by these bonds [145].

In the promoter region of a gene, cis-elements and transcription factors (TFs) interact, where they initiate gene transcription by assembling into the transcription initiation complex and then activating the RNA polymerase. [101]. Sixty-nine cis-elements were identified in *OsAPY*s and categorized into eight groups. They were essential in regulating various kinds of stresses and developmental stages, which indicates the engagement of *OsAPY*s in such situations. In plants, the evolution and expression of a character are determined by the gene’s chromosomal position. [107]. In this study, genes were unevenly distributed throughout the genome, and chromosomes 3, 7, 8, 10, 11, 12 contained all the *OsAPY* genes, and chromosome 11 contained the maximum number of *OsAPY* genes.

We discovered six distinct hormone-sensitive elements within the promoter region, including abscisic acid, gibberellin, auxin, salicylic acid, ethylene, and methyl-jasmonate responsive cis-elements. The involvement of *APY* in maintaining the level of auxin and ethylene [146] and in controlling the closure of stomata in drought condition induced by abscisic acid [147] was well documented. Moreover, there is ample evidence regarding the crosstalk between ethylene, gibberellin, and auxin [148,149,150]. These previous findings and the present study indicate the probable significance of *OsAPY*s in respective hormonal pathways.

miRNA-mediated regulation of genes has a vital function in controlling the environmental stimuli responsiveness of plants [38]. In this investigation, 103 miRNAs which target the *OsAPY* family members of rice were identified. Twenty-three miRNAs targeted more than one gene, but the rest were specific to only one gene. osa-miR5819 targeted four different genes. It implies that osa-miR5819 could have an important function in the expression of *OsAPY*s. Maximum of the miRNAs were found to be downregulated in many stresses and tissues, with exceptions of osa-miR159a.1, osa-miR5532, osa-miR408-3p and osa-miR3979-3p. Comparatively, the miRNAs showed reduced expression when subjected to various stresses; as plants adapt to growth conditions and environmental stressors, the miRNA-mediated control of *OsAPY* genes could have a decisive role.

As per the 3K SNP search database, the prevalence of nonsynonymous SAPs in *OsAPY*s is significantly greater for indica rice cultivars than other rice cultivars. *OsAPY1*, *5*, and *8* showed no SAPs in the studied varieties. Results indicate that the *OsAPY1*, *5*, and *8* are substantially conserved among the studied genotypes. *OsAPY2* contained the maximum number of SAPs which was 5 SAPs. The studied varieties are stress-responsive [38, 75] the presence of SAPs indicates their stress responsiveness might be related to these SAPs [151]. For fully comprehending their stress response basis, extensive research is required.

The involvement of *OsAPY*s in various tissues during the development and growth of rice was determined by an analysis of their expression patterns (Fig 11) using RNA-seq data. *OsAPY1- 4*, *9* showed higher levels of gene expression in inflorescence, implying their possible function in flower development. Earlier studies have proven the role of *APY* in flowers [25, 152]. *OsAPY2* and *4* had an abundant expression in anther, whereas *OsAPY1*, *3-5* showed elevated expression in the pistil, suggesting their potential roles in the germination of pollen, fertilization, and reproduction, respectively. The activity of *APY*s in pollen and pistils was reported in *Arabidopsis* [25, 32]. *OsAPY9* expression was found to be higher in the leaves and seedling stage, indicating that it plays a potential function in leaf development and early phases of plant growth. The involvement of *APY*s in leaves has previously been documented in *Medicago truncatula* [152] and *Arabidopsis* [25]. Another research on pea seedlings found that *APY*s were involved in early growth and development [153].

To further consider the dynamics of the *OsAPY*s in various biotic and abiotic stresses, their expression profiles via RNA-seq data were studied. The activity of *OsAPY*s was investigated in response to bacteria, fungus, and viruses, and *OsAPY1-5*, and *8* exhibited higher activation. *APY*s have been discovered to function in defensive and symbiotic relationships between plants and microorganisms [154]. *OsAPY1-5* demonstrated higher activity under blight-causing bacteria *Xanthomonas oryzae* pv. *oryzicola*. Both types of viral infections elevated the expression of *OsAPY2, 3* and *5*; under fungal infection, the expression of *OsAPY2*, *5*, and *8* increased. This discovery is in line with an earlier study that found the pea *APY* gene (*PsAPY1*) to be engaged in delivering protection against fungal infections [155]. The analysis of cis- elements showed that in the promoter region of *OsAPY*s, WRE3, WUN motifs, and W-box, that regulate biotic stresses in plants were present. The presence of such motifs corroborates our findings regarding the involvement of these genes in providing a defence under such stress conditions.

To identify the role of *OsAPY*s in abiotic stresses, the data retrieved from the RNA-seq database and the RT-qPCR analysis result were correlated. *OsAPY2* and *8* were essential genes concerning cold stress in both cases, suggesting their potential role under cold condition. In cold condition, *APY* involvement has been reported in *Populus euphratica* (*PeAPY2*) [134]. *OsAPY1*, *2*, and *5* showed increased activity under submergence stress suggesting their probable involvement in this condition. *OsAPY1-6* had upregulated expression under heat stress, suggesting these genes’ involvement in heat stress. Under salt stress, *OsAPY2* and *6* demonstrated elevated expression in both cases, indicating their role in helping the plant survive under salt stress. Similar results were found in wheat, where majority of the *APY* genes had higher expression after being exposed to salt for 12 hours in both roots and leaves [23].

*OsAPY5* showed higher expression under drought conditions indicating its role in helping plants cope with this condition, as the role of *APY* was previously reported in a study where the pea ectoapyrase was expressed in soybean and *Arabidopsis* and it resulted in better growth and tolerance towards drought conditions [156]. Presence of many abiotic stress responsive cis-elements support the notion concerning the role of *APY*s in abiotic stress conditions. Gene expression is one of the complicated biological processes, and it requires further research for elucidating mechanisms of the *OsAPY*s behind the regulation of several stresses. These findings indicate the function of *OsAPY*s under diverse biotic and abiotic stresses, and so this gene family could prove to be an important candidate for genetic engineering that can provide protection against various stress conditions.

## Conclusion

In this study, the rice *APY* gene family has been identified, characterized, and its expression profiling has been done extensively. Nine genes were identified, and they were found to be located on chromosome 3, 7, 8, 10, 11 and 12. Phylogenetic analysis grouped the nine identified *OsAPY* genes into three groups, and the identified cis-elements revealed their involvement in different stress conditions, hormonal regulation, and developmental stages. This ancient gene family evolved via segmental duplication and some SAPs were present in the stress-responsive varieties, which might contribute to their response to stresses. Four different categories of bonds were identified between the OsAPYs and ATP, suggesting strong docking interactions between them. miRNA analysis helped in understanding the function of miRNA in modulating *OsAPY* activity, while expression profiling via RNA-seq data unveiled the role of *OsAPY*s under various growth phases as well as stresses. The expression analysis using RT-qPCR data also confirmed the response of *OsAPY*s in various abiotic stress conditions, especially in heat stress. These findings would give the insight to broaden the understanding and knowledge regarding *APY*s in rice as well as provide foundation in future research regarding *OsAPY*s also in the genomic alterations towards the improving of rice and eventually developing stress-tolerant varieties.

## Acknowledgements

The authors are thankful to the Plant Genetic Engineering Lab, Department of Genetic Engineering and Biotechnology, Shahjalal University of Science and Technology, Sylhet for supporting to conduct this research.

## Supporting information

**S1 Fig. Apyrase Conserved Region (ACR) analysis of the OsAPYs.** The alignment was done using MEGA X and visualized via GeneDoc. The navy blue, pink, and aqua colors indicate the amino acids that are conserved 100%, 80%, and 60%, respectively, and the red-colored boxes depict the apyrase conserved region (ACR).

**S2 Fig. Ramachandran plot of the tertiary structures of OsAPYs.** The Ramachandran plot was generated via PROCHECK. Residues in the most favored, additional allowed, generously allowed, and disallowed regions are specified via red, yellow, pale yellow, and white colors, respectively.

**S3 Fig. ERRAT plot of the tertiary structures of OsAPYs.** ERRAT plot was generated via ERRAT. Yellow bars indicate the segment of the proteins which could be excluded at a 95% confidence level, and the red bars denote the ones at a 99% confidence level. The section with a lower error rate is marked by white bars.

**S4 Fig. Z-score plot of the tertiary structures of OsAPYs.** The Z-score plot was generated via ProSA-web. The light blue color indicates the Z-score of the proteins measured by X-ray crystallography, and the dark blue color indicates the Z-score of the proteins measured by nuclear magnetic resonance (NMR) spectroscopy. Black dots indicate the Z-score of each protein.

**S5 Fig. Energy plot of the tertiary structures of OsAPYs.** The energy plot was generated via ProSA-web. The dark green line represents the energy averaged across each fragment of 40 residues, and the light green line depicts the one across each fragment of 10 residues.

**S6 Fig. Ramachandran plot of the protein (OsAPYs)- ligand (ATP) complex structure.** The Ramachandran plot was generated via PROCHECK. Residues in the most favored, additional allowed, generously allowed, and disallowed regions are specified via red, yellow, pale yellow, and white colors, respectively.

**S7 Fig. ERRAT plot of the protein (OsAPYs)- ligand (ATP) complex structure.** ERRAT plot was generated via ERRAT. Yellow bars indicate the segment of the proteins which could be excluded at a 95% confidence level, and the red bars denote the ones at a 99% confidence level. The section with a lower error rate is marked by white bars.

**S1 Table. Description of the conserved motifs in OsAPYs.**

S2 Table. Duplicated *APY* genes and the probable dates of duplication blocks in rice.

S3 Table. The location of all the *OsAPY*s in different chromosome, their position, and orientation.

S4 Table. List of miRNAs targeting different *OsAPY*s and their mode of inhibition.

**S5 Table. The percentage of alpha-helix, beta-sheet, beta bridge, turn, coil, 310 helix, and the position of the transmembrane motif in the OsAPY members.**

**S6 Table. Validation score of OsAPY tertiary structure via different tools.**

**S7 Table. Docking score of the docked complexes generated via HDOCK server.**

**S8 Table: Number of bonds formed between OsAPYs and ATP according to PDBSum analysis.**

**S9 Table. Analysis of the interaction of OsAPY proteins with their ligand (ATP). S10 Table. List of primers used in RT-qPCR analysis.**

## References

1. Knowles AF. The GDA1_CD39 superfamily: NTPDases with diverse functions. Purinergic Signal. 2011 Mar;7(1):21–45.

2. Chiu TY, Lao J, Manalansan B, Loqué D, Roux SJ, Heazlewood JL. Biochemical characterization of *Arabidopsis* APYRASE family reveals their roles in regulating endomembrane NDP/NMP homoeostasis. Biochem J. 2015 Nov 15;472(1):43–54.

3. Thomas C, Rajagopal A, Windsor B, Dudler R, Lloyd A, Roux SJ. A Role for Ectophosphatase in Xenobiotic Resistance. Plant Cell. 2000 Apr;12(4):519–33.

4. Wu J, Steinebrunner I, Sun Y, Butterfield T, Torres J, Arnold D, et al. Apyrases (Nucleoside Triphosphate-Diphosphohydrolases) Play a Key Role in Growth Control in Arabidopsis. Plant Physiol. 2007 Jun;144(2):961–75.

5. Song CJ, Steinebrunner I, Wang X, Stout SC, Roux SJ. Extracellular ATP Induces the Accumulation of Superoxide via NADPH Oxidases in Arabidopsis. Plant Physiol. 2006 Apr;140(4):1222–32.

6. Kim SY, Sivaguru M, Stacey G. Extracellular ATP in Plants. Visualization, Localization, and Analysis of Physiological Significance in Growth and Signaling. Plant Physiol. 2006 Nov;142(3):984–92.

7. Wu SJ, Liu YS, Wu JY. The Signaling Role of Extracellular ATP and its Dependence on Ca^2+^ Flux in Elicitation of *Salvia miltiorrhiza* Hairy Root Cultures. Plant Cell Physiol. 2008 Apr;49(4):617–24.

8. Chivasa S, Ndimba BK, Simon WJ, Lindsey K, Slabas AR. Extracellular ATP Functions as an Endogenous External Metabolite Regulating Plant Cell Viability. The Plant Cell. 2005 Nov;17(11):3019–34.

9. Jeter CR, Tang W, Henaff E, Butterfield T, Roux SJ. Evidence of a novel cell signaling role for extracellular adenosine triphosphates and diphosphates in Arabidopsis. The Plant Cell. 2004 Oct;16(10):2652–64.

10. Kim SH, Yang SH, Kim TJ, Han JS, Suh JW. Hypertonic Stress Increased Extracellular ATP Levels and the Expression of Stress-Responsive Genes in *Arabidopsis thaliana* Seedlings. Biosci Biotechnol Biochem. 2009 Jun;73(6):1252–6.

11. Weerasinghe RR, Swanson SJ, Okada SF, Garrett MB, Kim SY, Stacey G, et al. Touch induces ATP release in Arabidopsis roots that is modulated by the heterotrimeric G- protein complex. FEBS Lett. 2009 Aug 6;583(15):2521–6.

12. Huang GT, Ma SL, Bai LP, Zhang L, Ma H, Jia P, et al. Signal transduction during cold, salt, and drought stresses in plants. Mol Biol Rep. 2012 Feb;39(2):969–87.

13. Zhu JK. Salt and Drought Stress Signal Transduction in Plants. Annu Rev Plant Biol. 2002;53:247–73.

14. Roux SJ, Steinebrunner I. Extracellular ATP: an unexpected role as a signaler in plants. Trends Plant Sci. 2007 Nov;12(11):522–527.

15. Tang W, Brady SR, Sun Y, Muday GK, Roux SJ. Extracellular ATP Inhibits Root Gravitropism at Concentrations That Inhibit Polar Auxin Transport. Plant Physiol. 2003 Jan;131(1):147–54.

16. Demidchik V, Shang Z, Shin R, Thompson E, Rubio L, Laohavisit A, et al. Plant extracellular ATP signalling by plasma membrane NADPH oxidase and Ca^2+^ channels. Plant J. 2009 Jun;58(6):903–13.

17. Tanaka K, Gilroy S, Jones AM, Stacey G. Extracellular ATP signaling in plants. Trends Cell Biol. 2010 Oct;20(10):601–8.

18. Clark G, Roux SJ. Role of Ca^2+^ in Mediating Plant Responses to Extracellular ATP and ADP. Int J Mol Sci.. 2018 Nov 14;19(11):3590.

19. Lim MH, Wu J, Yao J, Gallardo IF, Dugger JW, Webb LJ, et al. Apyrase Suppression Raises Extracellular ATP Levels and Induces Gene Expression and Cell Wall Changes Characteristic of Stress Responses. Plant Physiol. 2014 Apr;164(4):2054–67.

20. Sun J, Zhang CL, Deng SR, Lu CF, Shen X, Zhou XY, et al. An ATP signalling pathway in plant cells: extracellular ATP triggers programmed cell death in *Populus euphratica*. Plant Cell Environ. 2012 May;35(5):893–916.

21. Clark G, Brown KA, Tripathy MK, Roux SJ. Recent Advances Clarifying the Structure and Function of Plant Apyrases (Nucleoside Triphosphate Diphosphohydrolases). Int J Mol Sci. 2021 Mar 23;22(6):3283.

22. Zimmermann H, Zebisch M, Sträter N. Cellular function and molecular structure of ecto- nucleotidases. Purinergic Signal. 2012 Sep;8(3):437–502.

23. Liu W, Ni J, Shah FA, Ye K, Hu H, Wang Q, et al. Genome-wide identification, characterization and expression pattern analysis of APYRASE family members in response to abiotic and biotic stresses in wheat. PeerJ. 2019 Sep 11;7:e7622.

24. Clark GB, Morgan RO, Fernandez MP, Salmi ML, Roux SJ. Breakthroughs spotlighting roles for extracellular nucleotides and apyrases in stress responses and growth and development. Plant Sci. 2014 Aug 1;225:107–16.

25. Yang J. Functional analyses of Arabidopsis apyrases 3 through 7. In: Molecular, cell and developmental biology. Doctoral Dissertation. The University of Texas at Austin, Austin, TX. 2011. Available from: https://repositories.lib.utexas.edu/handle/2152/ETD-UT-2011-05-3056.

26. Leal DB, Streher CA, Neu TN, Bittencourt FP, Leal CA, da Silva JE, et al. Characterization of NTPDase (NTPDase1; ecto-apyrase; ecto-diphosphohydrolase; CD39; EC 3.6.1.5) activity in human lymphocytes. Biochim Biophys Acta. 2005 Jan 18;1721(1-3):9–15.

27. Handa M, Guidotti G. Purification and Cloning of a Soluble ATP-Diphosphohydrolase (Apyrase) from Potato Tubers (*Solanum tuberosum*). Biochem Biophys Res Commun. 1996 Jan 26;218(3):916–23.

28. Riewe D, Grosman L, Fernie AR, Wucke C, Geigenberger P. The Potato-Specific Apyrase Is Apoplastically Localized and Has Influence on Gene Expression, Growth, and Development. Plant Physiol. 2008 Jul;147(3):1092–109.

29. Day RB, McAlvin CB, Loh JT, Denny RL, Wood TC, Young ND, et al. Differential Expression of Two Soybean Apyrases, One of Which Is an Early Nodulin. Mol Plant Microbe Interact. 2000 Oct;13(10):1053–70.

30. Clark G, Torres J, Finlayson S, Guan X, Handley C, Lee J, et al. Apyrase (Nucleoside Triphosphate-Diphosphohydrolase) and Extracellular Nucleotides Regulate Cotton Fiber Elongation in Cultured Ovules. Plant Physiol. 2010 Feb;152(2):1073–83.

31. Steinebrunner I, Wu J, Sun Y, Corbett A, Roux SJ. Disruption of Apyrases Inhibits Pollen Germination in Arabidopsis. Plant Physiol. 2003 Apr;131(4):1638–47.

32. Yang J, Wu J, Romanovicz D, Clark G, Roux SJ. Co-regulation of exine wall patterning, pollen fertility and anther dehiscence by Arabidopsis apyrases 6 and 7. Plant Physiol Biochem. 2013 Aug;69:62–73.

33. Tan Y, Deng S, Sun Y, Zhao R, Jin X, Shen Z, et al. Functional analysis of *Populus euphratica PeAPY1* and *PePY2* in enhancing salt and drought tolerance. Genomics Appl Biol. 2014;33(4):860–8.

34. Alam I, Lee DG, Kim KH, Park CH, Sharmin SA, Lee H, et al. Proteome analysis of soybean roots under waterlogging stress at an early vegetative stage. J Biosci. 2010 Mar;35(1):49–62.

35. Shiraishi T. Suppression of Defense Response Related to Plant Cell Wall. Japan Agricultural Research Quarterly: Jpn. Agric. Res. Q. 2013;47(1):21–7.

36. Sivamaruthi BS, Kesika P, Chaiyasut C. Anthocyanins in Thai rice varieties: distribution and pharmacological significance. Int Food Res J. 2018 Sep 1;25(5):2024–32.

37. Van Nguyen N, Ferrero A. Meeting the challenges of global rice production. Paddy Water Environ. 2006 Mar;4(1):1–9.

38. Yadav SK, Santosh Kumar VV, Verma RK, Yadav P, Saroha A, Wankhede DP, et al. Genome-wide identification and characterization of ABA receptor *PYL* gene family in rice. BMC Genomics. 2020 Sep 30;21(1):676.

39. Nuruzzaman M, Manimekalai R, Sharoni AM, Satoh K, Kondoh H, Ooka H, et al. Genome-wide analysis of NAC transcription factor family in rice. Gene. 2010 Oct 1;465(1-2):30–44.

40. Tian C, Wan P, Sun S, Li J, Chen M. Genome-Wide Analysis of the GRAS Gene Family in Rice and *Arabidopsis*. Plant Mol Biol. 2004 Mar;54(4):519–32.

41. Katiyar A, Smita S, Lenka SK, Rajwanshi R, Chinnusamy V, Bansal KC. Genome-wide classification and expression analysis of *MYB* transcription factor families in rice and Arabidopsis. BMC Genom. 2012 Dec;13(1):1–9.

42. Ross CA, Liu Y, Shen QJ. The *WRKY* Gene Family in Rice (*Oryza sativa*). J Integr Plant Biol. 2007 Jun;49(6):827–42.

43. Zhao L, Hu Y, Chong K, Wang T. *ARAG1*, an ABA-responsive DREB gene, plays a role in seed germination and drought tolerance of rice. Ann Bot. 2010 Mar;105(3):401–9.

44. Jung KH, Ko HJ, Nguyen MX, Kim SR, Ronald P, An G. Genome-wide identification and analysis of early heat stress responsive genes in rice. J Plant Biol. 2012 Dec;55(6):458–68.

45. Shabala S. Regulation of Potassium Transport in Leaves: from Molecular to Tissue Level. Ann Bot. 2003 Nov;92(5):627–34.

46. Yamaguchi-Shinozaki K, Shinozaki K. Organization of *cis*-acting regulatory elements in osmotic- and cold-stress-responsive promoters. Trends Plant Sci. 2005 Feb;10(2):88–94.

47. Amin US, Biswas S, Elias SM, Razzaque S, Haque T, Malo R, et al. Enhanced Salt Tolerance Conferred by the Complete 2.3 kb cDNA of the Rice Vacuolar Na^+^/H^+^ Antiporter Gene Compared to 1.9 kb Coding Region with 5′ UTR in Transgenic Lines of Rice. Front Plant Sci. 2016 Jan 25;7:14.

48. Chen JQ, Meng XP, Zhang Y, Xia M, Wang XP. Over-expression of OsDREB genes lead to enhanced drought tolerance in rice. Biotechnol Lett. 2008 Dec;30(12):2191–8.

49. Xu K, Xu X, Fukao T, Canlas P, Maghirang-Rodriguez R, Heuer S, et al. *Sub1A* is an ethylene-response-factor-like gene that confers submergence tolerance to rice. Nature. 2006 Aug 10;442(7103):705–8.

50. Huala E, Dickerman AW, Garcia-Hernandez M, Weems D, Reiser L, LaFond F, et al. The Arabidopsis Information Resource (TAIR): a comprehensive database and web- based information retrieval, analysis, and visualization system for a model plant. Nucleic Acids Res. 2001 Jan 1;29(1):102–5.

51. Goodstein DM, Shu S, Howson R, Neupane R, Hayes RD, Fazo J, et al. Phytozome: a comparative platform for green plant genomics. Nucleic Acids Res. 2012 Jan;40(D1):D1178–86.

52. Kawahara Y, de la Bastide M, Hamilton JP, Kanamori H, McCombie WR, Ouyang S, et al. Improvement of the *Oryza sativa* Nipponbare reference genome using next generation sequence and optical map data. Rice. 2013 Feb 6;6(1):4.

53. Hunter S, Apweiler R, Attwood TK, Bairoch A, Bateman A, Binns D, et al. InterPro: the integrative protein signature database. Nucleic Acids Res. 2009 Jan;37(suppl_1):D211–5.

54. Kumar S, Stecher G, Li M, Knyaz C, Tamura K. MEGA X: Molecular Evolutionary Genetics Analysis across Computing Platforms. Mol Biol Evol. 2018 Jun 1;35(6):1547–1549.

55. Nicholas KB. GeneDoc: analysis and visualization of genetic variation. Embnew news. 1997;4:14.

56. Wilkins MR, Gasteiger E, Bairoch A, Sanchez JC, Williams KL, Appel RD, et al. Protein Identification and Analysis Tools on the ExPASy Server. Methods Mol Biol. 1999;112:531–52.

57. Horton P, Park KJ, Obayashi T, Fujita N, Harada H, Adams-Collier CJ, et al. WoLF PSORT: protein localization predictor. Nucleic Acids Res. 2007 Jul;35(suppl_2):W585–7.

58. Yu CS, Chen YC, Lu CH, Hwang JK. Prediction of protein subcellular localization. Proteins. 2006 Aug 15;64(3):643–51.

59. Bailey TL, Boden M, Buske FA, Frith M, Grant CE, Clementi L, et al. MEME SUITE: tools for motif discovery and searching. Nucleic Acids Res. 2009 Jul 1;37(suppl_2):W202–8.

60. El-Gebali S, Mistry J, Bateman A, Eddy SR, Luciani A, Potter SC, et al. The Pfam protein families database in 2019. Nucleic Acids Res. 2019 Jan 8;47(D1):D427–D432.

61. Hu B, Jin J, Guo AY, Zhang H, Luo J, Gao G. GSDS 2.0: an upgraded gene feature visualization server. Bioinformatics. 2015 Apr 15;31(8):1296–7.

62. Bolser D, Staines DM, Pritchard E, Kersey P. Ensembl Plants: Integrating Tools for Visualizing, Mining, and Analyzing Plant Genomics Data. Methods Mol Biol. 2016;1374:115–40.

63. Guindon S, Gascuel O. A Simple, Fast, and Accurate Algorithm to Estimate Large Phylogenies by Maximum Likelihood. Syst Biol. 2003 Oct;52(5):696–704.

64. Letunic I, Bork P. Interactive tree of life (iTOL) v3: an online tool for the display and annotation of phylogenetic and other trees. Nucleic Acids Res. 2016 Jul 8;44(W1):W242–5.

65. Lee TH, Tang H, Wang X, Paterson AH. PGDD: a database of gene and genome duplication in plants. Nucleic Acids Res. 2013 Jan;41(D1):D1152–8.

66. Koch MA, Haubold B, Mitchell-Olds T. Comparative Evolutionary Analysis of Chalcone Synthase and Alcohol Dehydrogenase Loci in *Arabidopsis*, *Arabis*, and Related Genera (Brassicaceae). Mol Biol Evol. 2000 Oct;17(10):1483–98.

67. Chen C, Chen H, Zhang Y, Thomas HR, Frank MH, He Y, et al. TBtools: An Integrative Toolkit Developed for Interactive Analyses of Big Biological Data. Mol Plant. 2020 Aug 3;13(8):1194–202.

68. Lescot M, Déhais P, Thijs G, Marchal K, Moreau Y, Van de Peer Y, et al. PlantCARE, a database of plant *cis*-acting regulatory elements and a portal to tools for *in silico* analysis of promoter sequences. Nucleic Acids Res. 2002 Jan 1;30(1):325–7.

69. Voorrips RE. MapChart: Software for the Graphical Presentation of Linkage Maps and QTLs. J Hered. 2002 Jan-Feb;93(1):77–8.

70. Dai X, Zhao PX. psRNATarget: a plant small RNA target analysis server. Nucleic Acids Res. 2011 Jul;39(suppl_2):W155–9.

71. Kozomara A, Griffiths-Jones S. miRBase: annotating high confidence microRNAs using deep sequencing data. Nucleic Acids Res. 2014 Jan;42(D1):D68–73.

72. Shannon P, Markiel A, Ozier O, Baliga NS, Wang JT, Ramage D, et al. Cytoscape: A Software Environment for Integrated Models of Biomolecular Interaction Networks. Genome Res. 2003 Nov;13(11):2498–504.

73. Gurjar AK, Panwar AS, Gupta R, Mantri SS. PmiRExAt: plant miRNA expression atlas database and web applications. Database (Oxford). 2016 Apr 13;2016:baw060.

74. 74. Motulsky HJ. Analyzing data with graphpad prism. GraphPad Software. Inc., San Diego, CA. 1999.

75. Kaur V, Yadav SK, Wankhede DP, Pulivendula P, Kumar A, Chinnusamy V. Cloning and characterization of a gene encoding MIZ1, a domain of unknown function protein and its role in salt and drought stress in rice. Protoplasma. 2020 Mar;257(2):475–487.

76. Mansueto L, Fuentes RR, Borja FN, Detras J, Abriol-Santos JM, Chebotarov D, et al. Rice SNP-seek database update: new SNPs, indels, and queries. Nucleic Acids Res. 2017 Jan 4;45(D1):D1075–D1081.

77. Heinig M, Frishman D. STRIDE: a web server for secondary structure assignment from known atomic coordinates of proteins. Nucleic Acids Res. 2004 Jul 1;32(suppl_2):W500–2.

78. Krogh A, Larsson B, von Heijne G, Sonnhammer EL. Predicting transmembrane protein topology with a hidden Markov model: application to complete genomes. J Mol Biol. 2001 Jan 19;305(3):567–80.

79. Kim DE, Chivian D, Baker D. Protein structure prediction and analysis using the Robetta server. Nucleic Acids Res. 2004 Jul 1;32(suppl_2):W526–31..

80. Ko J, Park H, Heo L, Seok C. GalaxyWEB server for protein structure prediction and refinement. Nucleic Acids Res. 2012 Jul;40(W1):W294–7.

81. Guex N, Peitsch MC. SWISS-MODEL and the Swiss-PdbViewer: An environment for comparative protein modeling. Electrophoresis. 1997 Dec;18(15):2714–23.

82. Laskowski RA, Rullmannn JA, MacArthur MW, Kaptein R, Thornton JM. AQUA and PROCHECK-NMR: Programs for checking the quality of protein structures solved by NMR. J Biomol NMR. 1996 Dec;8(4):477–86.

83. Colovos C, Yeates TO. Verification of protein structures: Patterns of nonbonded atomic interactions. Protein Sci. 1993 Sep;2(9):1511–9.

84. Wiederstein M, Sippl MJ. ProSA-web: interactive web service for the recognition of errors in three-dimensional structures of proteins. Nucleic Acids Res. 2007 Jul;35(suppl_2):W407–10.

85. Systemes, D., BIOVIA, discovery studio modeling environment. Release 4.5. Dassault Systemes: San Diego, CA. 2015.

86. Yan Y, Zhang D, Zhou P, Li B, Huang SY. HDOCK: a web server for protein–protein and protein–DNA/RNA docking based on a hybrid strategy. Nucleic Acids Res. 2017 Jul 3;45(W1):W365–W373.

87. Laskowski RA. PDBsum: summaries and analyses of PDB structures. Nucleic Acids Res. 2001 Jan 1;29(1):221–2.

88. Hruz T, Laule O, Szabo G, Wessendorp F, Bleuler S, Oertle L, et al. Genevestigator V3: A Reference Expression Database for the Meta-Analysis of Transcriptomes. Adv Bioinformatics. 2008;2008:420747.

89. Untergasser A, Cutcutache I, Koressaar T, Ye J, Faircloth BC, Remm M, et al. Primer3--new capabilities and interfaces. Nucleic Acids Res. 2012 Aug;40(15):e115.

90. Jain M, Nijhawan A, Tyagi AK, Khurana JP. Validation of housekeeping genes as internal control for studying gene expression in rice by quantitative real-time PCR. Biochem Biophys Res Commun. 2006 Jun 30;345(2):646–51.

91. Livak KJ, Schmittgen TD. Analysis of Relative Gene Expression Data Using Real-Time Quantitative PCR and the 2^−ΔΔCT^ Method. Methods. 2001 Dec;25(4):402–8.

92. Chang KY, Yang JR. Analysis and Prediction of Highly Effective Antiviral Peptides Based on Random Forests. PLoS One. 2013 Aug 5;8(8):e70166.

93. Liu Z, Liu Y, Coulter JA, Shen B, Li Y, Li C, et al. The WD40 Gene Family in Potato (*Solanum Tuberosum* L.): Genome-Wide Analysis and Identification of Anthocyanin and Drought-Related WD40s. Agronomy. 2020 Mar 15;10(3):401.

94. Hattori T, Totsuka M, Hobo T, Kagaya Y, Yamamoto-Toyoda A. Experimentally determined sequence requirement of ACGT-containing abscisic acid response element. Plant Cell Physiol. 2002 Jan;43(1):136–40.

95. Liu X, Sun Z, Dong W, Wang Z, Zhang L. Expansion and Functional Divergence of the SHORT VEGETATIVE PHASE (SVP) Genes in Eudicots. Genome Biol Evol. 2018 Nov 1;10(11):3026–3037.

96. Gubler F, Jacobsen JV. Gibberellin-responsive elements in the promoter of a barley high-pI alpha-amylase gene. Plant Cell. 1992 Nov;4(11):1435–41.

97. Ogawa M, Hanada A, Yamauchi Y, Kuwahara A, Kamiya Y, Yamaguchi S. Gibberellin biosynthesis and response during Arabidopsis seed germination. Plant Cell. 2003 Jul;15(7):1591–604.

98. Heis MD, Ditmer EM, de Oliveira LA, Frazzon AP, Margis R, Frazzon J. Differential expression of cysteine desulfurases in soybean. BMC Plant Biol. 2011 Nov 18;11:166.

99. Yin J, Li X, Zhan Y, Li Y, Qu Z, Sun L, et al. Cloning and expression of BpMYC4 and BpbHLH9 genes and the role of BpbHLH9 in triterpenoid synthesis in birch. BMC Plant Biol. 2017 Nov 21;17(1):214.

100. Shokri-Gharelo R, Bandehagh A, Mahmoudi B, Moti-Noparvar P. In silico study of cis-acting elements revealing the plastid gene involved in oxidative phosphorylation are responsive to abiotic stresses. Acta Biol. Szeged. 2017 Jan 1;61(2):179–88.

101. Sutoh K, Yamauchi D. Two cis-acting elements necessary and sufficient for gibberellin-upregulated proteinase expression in rice seeds. Plant J. 2003 Jun;34(5):635–45.

102. Kim SR, Kim Y, An G. Identification of methyl jasmonate and salicylic acid response elements from the nopaline synthase (nos) promoter. Plant Physiol. 1993 Sep;103(1):97–103.

103. Gong XX, Yan BY, Tan YR, Gao X, Wang D, Zhang H, et al. Identification of cis- regulatory regions responsible for developmental and hormonal regulation of HbHMGS1 in transgenic *Arabidopsis thaliana*. Biotechnol Lett. 2019 Sep;41(8-9):1077–1091.

104. Li N, Wei S, Chen J, Yang F, Kong L, Chen C, et al. OsASR2 regulates the expression of a defence-related gene, Os2H16, by targeting the GT-1 cis-element. Plant Biotechnol J. 2018 Mar;16(3):771–783.

105. Zuckerman GR, Prakash C, Askin MP, Lewis BS. AGA technical review on the evaluation and management of occult and obscure gastrointestinal bleeding. Gastroenterology. 2000 Jan;118(1):201–21.

106. Arefin S, Bhuiyan MF, Akther J, Prodhan SH, Hoque H. The dynamics of cis-regulatory elements in promoter regions of tomato sucrose transporter genes. Int J Veg Sci. 2021 Mar 4;27(2):167–86.

107. Piechulla B, Merforth N, Rudolph B. Identification of tomato Lhc promoter regions necessary for circadian expression. Plant Mol Biol. 1998 Nov 1;38(4):655–62.

108. Ezcurra I, Ellerström M, Wycliffe P, Stålberg K, Rask L. Interaction between composite elements in the napA promoter: both the B-box ABA-responsive complex and the RY/G complex are necessary for seed-specific expression. Plant Mol Biol. 1999 Jul;40(4):699–709.

109. Jin XQ, Chen ZW, Tan RH, Zhao SJ, Hu ZB. Isolation and functional analysis of 4- coumarate: coenzyme A ligase gene promoters from *Salvia miltiorrhiza*. Biol Plant. 2012 Jun;56(2):261–8.

110. Wang X, Zhang H, Shao LY, Yan X, Peng H, Ouyang JX, et al. Expression and function analysis of a rice OsHSP40 gene under salt stress. Genes Genomics. 2019 Feb;41(2):175–182.

111. Hou F, Du T, Qin Z, Xu T, Li A, Dong S, et al. Genome-wide in silico identification and expression analysis of beta-galactosidase family members in sweetpotato [*Ipomoea batatas* (L.) Lam]. BMC Genom. 2021 Dec;22(1):1–3.

112. Kaur A, Pati PK, Pati AM, Nagpal AK. In-silico analysis of cis-acting regulatory elements of pathogenesis-related proteins of *Arabidopsis thaliana* and *Oryza sativa*. PLoS One. 2017 Sep 14;12(9):e0184523.

113. Lois R, Dietrich A, Hahlbrock K, Schulz W. A phenylalanine ammonia-lyase gene from parsley: structure, regulation and identification of elicitor and light responsive cis-acting elements. EMBO J. 1989 Jun;8(6):1641–8.

114. López-Ochoa L, Acevedo-Hernández G, Martínez-Hernández A, Argüello-Astorga G, Herrera-Estrella L. Structural relationships between diverse cis-acting elements are critical for the functional properties of a rbcS minimal light regulatory unit. J Exp Bot. 2007;58(15-16):4397–406.

115. Liu L, Zhang X, Chen F, Mahi AA, Wu X, Chen Q, et al. Analysis of promoter activity reveals that *GmFTL2* expression differs from that of the known *Flowering Locus T* genes in soybean. Crop J. 2017 Oct 1;5(5):438–48.

116. Le Gourrierec J, Li YF, Zhou DX. Transcriptional activation by Arabidopsis GT-1 may be through interaction with TFIIA-TBP-TATA complex. Plant J. 1999 Jun;18(6):663–8.

117. De Silva WS, Perera MM, Perera KL, Wickramasuriya AM, Jayasekera GA. In silico analysis of osr40c1 promoter sequence isolated from Indica variety Pokkali. Rice Sci. 2017 Jul 1;24(4):228–34.

118. Hiratsuka K, Chua NH. Light regulated transcription in higher plants. J Plant Res. 1997 Mar;110(1):131–9.

119. Prabu G, Prasad DT. Functional characterization of sugarcane MYB transcription factor gene promoter (PScMYBAS1) in response to abiotic stresses and hormones. Plant Cell Rep. 2012 Apr;31(4):661–9.

120. Hazra A, Dasgupta N, Sengupta C, Das S. MIPS: Functional dynamics in evolutionary pathways of plant kingdom. Genomics. 2019 Dec;111(6):1929–1945.

121. Haralampidis K, Milioni D, Rigas S, Hatzopoulos P. Combinatorial interaction of cis elements specifies the expression of the Arabidopsis AtHsp90-1 gene. Plant Physiol. 2002 Jul;129(3):1138–49.

122. Wu A, Hao P, Wei H, Sun H, Cheng S, Chen P, et al. Genome-Wide Identification and Characterization of Glycosyltransferase Family 47 in Cotton. Front Genet. 2019 Sep 11;10:824.

123. Chen W, Provart NJ, Glazebrook J, Katagiri F, Chang HS, Eulgem T, et al. Expression profile matrix of Arabidopsis transcription factor genes suggests their putative functions in response to environmental stresses. Plant Cell. 2002 Mar;14(3):559–74.

124. Liu J, Wang F, Yu G, Zhang X, Jia C, Qin J, et al. Functional Analysis of the Maize C-Repeat/DRE Motif-Binding Transcription Factor CBF3 Promoter in Response to Abiotic Stress. Int J Mol Sci. 2015 May 28;16(6):12131–46.

125. Dubouzet JG, Sakuma Y, Ito Y, Kasuga M, Dubouzet EG, Miura S, Seki M, Shinozaki K, Yamaguchi-Shinozaki K. OsDREB genes in rice, *Oryza sativa* L., encode transcription activators that function in drought-, high-salt- and cold-responsive gene expression. Plant J. 2003 Feb;33(4):751–63.

126. Abe H, Urao T, Ito T, Seki M, Shinozaki K, Yamaguchi-Shinozaki K. Arabidopsis AtMYC2 (bHLH) and AtMYB2 (MYB) function as transcriptional activators in abscisic acid signaling. Plant Cell. 2003 Jan;15(1):63–78.

127. Li Z, Li G, Cai M, Priyadarshani SVGN, Aslam M, Zhou Q, et al. Genome-Wide Analysis of the YABBY Transcription Factor Family in Pineapple and Functional Identification of *AcYABBY4* Involvement in Salt Stress. Int J Mol Sci. 2019 Nov 22;20(23):5863.

128. Shinozaki K, Yamaguchi-Shinozaki K. Gene Expression and Signal Transduction in Water-Stress Response. Plant Physiol. 1997 Oct;115(2):327–334.

129. Pandey S, Subramanaym Reddy C, Yaqoob U, Negi Y, Arora S. Insilico analysis of cis acting regulatory elements CAREs in upstream regions of ascorbate glutathione pathway genes from *Oryza sativa*. Biochem Physiol. 2015;4(2).

130. Jiu S, Haider MS, Kurjogi MM, Zhang K, Zhu X, Fang J. Genome-wide Characterization and Expression Analysis of Sugar Transporter Family Genes in Woodland Strawberry. Plant Genome. 2018 Nov;11(3).

131. Tarasov KV, Tarasova YS, Tam WL, Riordon DR, Elliott ST, Kania G, et al. B-MYB is essential for normal cell cycle progression and chromosomal stability of embryonic stem cells. PLoS One. 2008 Jun 25;3(6):e2478.

132. Islam R, Rahaman M, Hoque H, Hasan N, Prodhan SH, Ruhama A, et al. Computational and structural based approach to identify malignant nonsynonymous single nucleotide polymorphisms associated with CDK4 gene. PLoS One. 2021 Nov 4;16(11):e0259691.

133. Muthukumaran J, Manivel P, Kannan M, Jeyakanthan J, Krishna R. A framework for classification of antifreeze proteins in over wintering plants based on their sequence and structural features. J. Bioinform. Seq. Anal. 2011 May;3:70–88.

134. Deng S, Sun J, Zhao R, Ding M, Zhang Y, Sun Y, et al. Populus euphratica APYRASE2 Enhances Cold Tolerance by Modulating Vesicular Trafficking and Extracellular ATP in Arabidopsis Plants. Plant Physiol. 2015 Sep;169(1):530–48.

135. Schneider P, Asch F. Rice production and food security in Asian Mega deltas—A review on characteristics, vulnerabilities and agricultural adaptation options to cope with climate change. J Agron Crop Sci. 2020 Aug;206(4):491–503.

136. Govindarajulu M, Kim SY, Libault M, Berg RH, Tanaka K, Stacey G, et al. GS52 Ecto-Apyrase Plays a Critical Role during Soybean Nodulation. Plant Physiol. 2009 Feb;149(2):994–1004.

137. Pietrowska-Borek M, Dobrogojski J, Sobieszczuk-Nowicka E, Borek S. New Insight into Plant Signaling: Extracellular ATP and Uncommon Nucleotides. Cells. 2020 Feb 2;9(2):345.

138. Huang Z, Duan W, Song X, Tang J, Wu P, Zhang B, et al. Retention, Molecular Evolution, and Expression Divergence of the Auxin/Indole Acetic Acid and Auxin Response Factor Gene Families in Brassica Rapa Shed Light on Their Evolution Patterns in Plants. Genome Biol Evol. 2016 Feb 1;8(2):302–16.

139. Wang Q, Xu X, Cao X, Hu T, Xia D, Zhu J, et al. Identification, Classification, and Expression Analysis of the Triacylglycerol Lipase (TGL) Gene Family Related to Abiotic Stresses in Tomato. Int J Mol Sci. 2021 Jan 30;22(3):1387.

140. Liu Q, Sun C, Han J, Li L, Wang K, Wang Y, et al. Identification, characterization and functional differentiation of the NAC gene family and its roles in response to cold stress in ginseng, Panax ginseng C.A. Meyer. PLoS One. 2020 Jun 11;15(6):e0234423.

141. Koralewski TE, Krutovsky KV. Evolution of Exon-Intron Structure and Alternative Splicing. PLoS One. 2011 Mar 25;6(3):e18055.

142. Deshmukh RK, Sonah H, Singh NK. Intron gain, a dominant evolutionary process supporting high levels of gene expression in rice. J Plant Biochem Biotechnol. 2016 Apr;25(2):142–6.

143. Nagar P, Kumar A, Jain M, Kumari S, Mustafiz A. Genome-wide analysis and transcript profiling of PSKR gene family members in *Oryza sativa*. PLoS One. 2020 Jul 23;15(7):e0236349.

144. MacIsaac KD, Fraenkel E. Practical Strategies for Discovering Regulatory DNA Sequence Motifs. PLoS Comput Biol. 2006 Apr;2(4):e36.

145. Elengoe A, Naser MA, Hamdan S. Modeling and Docking Studies on Novel Mutants (K71L and T204V) of the ATPase Domain of Human Heat Shock 70 kDa Protein 1. Int J Mol Sci. 2014 Apr 22;15(4):6797–814.

146. Liu X, Wu J, Clark G, Lundy S, Lim M, Arnold D, et al. Role for Apyrases in Polar Auxin Transport in Arabidopsis. Plant Physiol. 2012 Dec;160(4):1985–95.

147. Zhang Y, Sun Y, Liu X, Deng J, Yao J, Zhang Y, et al. Populus euphratica Apyrases Increase Drought Tolerance by Modulating Stomatal Aperture in Arabidopsis. Int J Mol Sci. 2021 Sep 13;22(18):9892.

148. Smalle J, Haegman M, Kurepa J, Van Montagu M, Straeten DV. Ethylene can stimulate Arabidopsis hypocotyl elongation in the light. Proc Natl Acad Sci U S A. 1997 Mar 18;94(6):2756–61.

149. Saibo NJ, Vriezen WH, Beemster GT, Van Der Straeten D. Growth and stomata development of Arabidopsis hypocotyls are controlled by gibberellins and modulated by ethylene and auxins. Plant J. 2003 Mar;33(6):989–1000.

150. Vandenbussche F, Smalle J, Le J, Saibo NJ, De Paepe A, Chaerle L, et al. The Arabidopsis Mutant alh1 Illustrates a Cross Talk between Ethylene and Auxin. Plant Physiol. 2003 Mar;131(3):1228–38.

151. Morgil H, Gercek YC, Tulum I. Single nucleotide polymorphisms (SNPs) in plant genetics and breeding. The Recent Topics in Genetic Polymorphisms: IntechOpen; 2020.

152. Navarro-Gochicoa MT, Camut S, Niebel A, Cullimore JV. Expression of the Apyrase-Like APY1 Genes in Roots of Medicago truncatula Is Induced Rapidly and Transiently by Stress and Not by Sinorhizobium meliloti or Nod Factors. Plant Physiol. 2003 Mar;131(3):1124–36.

153. Yoneda M, Davies E, Morita EH, Abe S. Immunohistochemical localization of apyrase during initial differentiation and germination of pea seeds. Planta. 2009 Dec;231(1):47–56.

154. Tanaka K, Tóth K, Stacey G. Role of ectoapyrases in nodulation. Biological Nitrogen Fixation. 2015 Aug 3:517–24.

155. Toyoda K, Yasunaga E, Niwa M, Ohwatari Y, Nakashima A, Inagaki Y, et al. H2O2 production by copper amine oxidase, a component of the ecto-apyrase (ATPase)- containing protein complex(es) in the pea cell wall, is regulated by an elicitor and a suppressor from Mycosphaerella pinodes. J Gen Plant Pathol. 2012 Sep;78(5):311–5.

156. Veerappa R, Slocum RD, Siegenthaler A, Wang J, Clark G, Roux SJ. Ectopic expression of a pea apyrase enhances root system architecture and drought survival in Arabidopsis and soybean. Plant Cell Environ. 2019 Jan;42(1):337–353.

